# RosENet: Improving binding affinity prediction by leveraging molecular mechanics energies with a 3D Convolutional Neural Network

**DOI:** 10.1101/2020.05.12.090191

**Authors:** Hussein Hassan-Harrirou, Ce Zhang, Thomas Lemmin

**Affiliations:** DS3Lab, System Group, Department of Computer Sciences, ETH Zurich, CH-8092 Zurich, Switzerland; Institute of Medical Virology, University of Zurich (UZH), CH-8057 Zurich, Switzerland

**Author notes:** Corresponding Author Thomas Lemmin.

**Keywords:** Convolutional Neural Network, binding affinity, molecular mechanics energies, drug discovery

## Abstract

The worldwide increase and proliferation of drug resistant microbes, coupled with the lag in new drug development represents a major threat to human health. In order to reduce the time and cost for exploring the chemical search space, drug discovery increasingly relies on computational biology approaches. One key step in these approaches is the need for the rapid and accurate prediction of the binding affinity for potential leads.

Here, we present RosENet (**Ros**etta **E**nergy Neural **Net**work), a three-dimensional (3D) Convolutional Neural Network (CNN), which combines voxelized molecular mechanics energies and molecular descriptors for predicting the absolute binding affinity of protein – ligand complexes. By leveraging the physico-chemical properties captured by the molecular force field, our model achieved a Root Mean Square Error (RMSE) of 1.26 on the PDBBind v2016 *core set*. We also explored some limitations and the robustness of the PDBBind dataset and our approach, on nearly 500 structures, including structures determined by Nuclear Magnetic Resonance and virtual screening experiments. Our study demonstrated that molecular mechanics energies can be voxelized and used to help improve the predictive power of the CNNs. In the future, our framework can be extended to features extracted from other biophysical and biochemical models, such as molecular dynamics simulations.

**Availability:** https://github.com/DS3Lab/RosENet

## INTRODUCTION

The alarming worldwide increase in drug resistant microbes is rapidly challenging and in certain cases, defeating the effectiveness of existing antibiotics, thus representing a serious threat to human health. The chemical search space for the discovery of small molecule therapeutics is immense, estimated at 10^66^ molecules. A brute force approach is thus not realistic. Advances in computational biology have proven to be valuable tools for more efficiently exploring this chemical search space and have led to the discovery of several leads for high affinity drugs. In addition, these computational methods permit significantly reducing the cost and workload for drug discovery.

One critical step in the drug design process is the scoring and ranking of the predicted drug – target interactions. Most methods aim to predict the binding affinity of the complex that represents the binding free energy in the target-ligand interactions. Experimentally, the binding affinity is commonly measured and reported by the equilibrium dissociation constant (K_D_), the inhibition constant (K_I_) or half maximal inhibitory concentration (IC_50_). The direct estimation of the free energy of the binding can be determined computationally with biased molecular simulations, such as umbrella sampling, thermodynamic integration and free energy perturbation. Although these methods have achieved in some cases very accurate estimates of the binding affinity (error smaller than 0.5 kcal/mol), they are extremely slow and compute intensive. Therefore, they are not adapted for large scale screens.

To overcome this problem, a variety of classical machine-learning algorithms have been applied, e.g., linear regressions;^1,2^ kernel ridge regression;^3,4^ support vector machines;^2,5^ Gaussian processes;^1^ random forests.^3,5,6^ RF-Score, one of the best performing models, is based on a random forest, that relies on 42 molecular descriptors extracted from AutoDock Vina.^7^ RF-Score reported a Root Mean Square Error (RMSE) of 1.51 for pK_D_ prediction on the PDBBind test set (v2007 *core set*). Its RMSE further decreased to 1.39, when trained with the larger 2016 version of PDBBind.^8,9^ AGL-Score utilizes a graph representation of the complexes to obtain statistical features of the adjacency and laplacian matrices of the graph.^10^ These statistics are then used as features in a gradient boosted decision tree. AGL-Score obtained an RMSE of 1.27 on the PDBBind v2016 *core set*.

Recently, Deep Learning approaches have allowed major breakthroughs in several fields of research and are being increasingly applied to structural biology and computational chemistry. Deep learning is a powerful framework built around neural networks, a class of algorithms inspired by the nervous system. These methods can learn more complex representations and automatically extract the features relevant to a problem.^11–13^ In particular, various Convolutional Neural Network (CNN) architectures have been used to predict the binding affinity. These include AtomNet,^14^ Atomic Convolutional Neural Network,^15^ TopologyNet,^16^ DeepDTA,^17^ DeepMHC,^18^ KDeep^19^, OnionNet^12^ and Pafnucy.^20^ These methods mainly differ by their feature extraction and embedding. For example, DeepDTA uses textual representations of protein and ligand for predicting pK_D_ values equal to or greater than 9 and achieves a Mean Square Error (MSE) of 0.261 on a split of the Davis dataset^21^ and 0.194 on a split of the KIBA dataset.^22^ The Atomic Convolutional Neural Network (ACNN) estimates the change in energy of the protein – ligand complex with an intermediate representation of pairwise atom distances and atom types obtained by custom convolution. ACNN reports an mean absolute error (MAE) of 0.77 kcal/mol on PDBBind’s *refined set*. TopologyNet uses topological fingerprints called Betti numbers with convolutions and predicts the pK_D_ with an RMSE of 1.37 on the PDBBind *core set*. OnionNet represents the pairwise interactions between 8 types of atoms with respect to a set of 60 distances. The resulting features were input to a two-dimensional (2D) convolutional neural network and obtained an RMSE of 1.28 on the PDBBind v2016 *core set*.

KDeep, Pafnucy and AtomNet employ a three dimensional (3D) image-like representation approach, where the attributes of the protein and ligand atoms are distributed on a 3D grid. AtomNet describes the protein – ligand complex with a combination of attributes ranging from atom types to complex interaction fingerprints and obtains an Area Under the ROC Curve (AUC) for virtual screening greater than 0.9 for nearly two-thirds of the targets in the DUD-E benchmark.^23^ Pafnucy employs 19 different atomic features, and reports an RMSE of 1.42 on the PDBBind v2016 *core set*. Finally, KDeep uses eight molecular descriptors defined by AutoDock Vina software and obtains an RMSE of 1.27 on the PDBBind v2016 *core set*. When tested on other datasets, KDeep was shown to be sensitive to the specific proteins.

Here, we present RosENet (**Ros**etta **E**nergy **Net**work), a 3D Convolutional Neural Network (CNN), which combines molecular mechanics energies (computed with the Rosetta force field^24^) with molecular descriptors. Molecular mechanics rely on a physical model for describing the interaction between atoms and have proven to be valuable methods for understanding the function and dynamics in biomolecular systems. They are therefore at the core of major computational tools for the modeling and design of biomolecules. We hypothesized that the high non-linearity and complexity of molecular mechanics energies could benefit from the use of deep neural network architectures. Since residual networks have the capacity to learn variations between the input and output, they would be ideal for capturing the binding differential of energy.

In this study, the molecular energies and descriptors were embedded onto a 3D grid and the CNN was based on the ResNet architecture.^25^ We tested RosENet on a set of 446 diverse protein – ligand complexes. In addition, we investigated the robustness RoseENet on virtual screening experiments and compared it with two recent CNN models, OnionNet^12^ and Pafnucy.^20^ The code, processed data and all generated datasets needed to reproduce our results are freely accessible on our GitHub repository.

## RESULTS

The data pre-processing, training and testing of RosENet was implemented as a modularized pipeline (**Figure 1**). In total, six different structural datasets were used for the training, validation and testing of RosENet. Each protein – ligand complex was first uniformized by renaming the protein, metallic, water and ligand chain identifiers and substituting non-standard residues with their standard counterparts. A minimization with the Rosetta software^26^ was then performed, where the ligand was randomly relaxed twenty times and the structure with the lowest total energy was saved. Next, the molecular energies and descriptors were voxelized onto a 25 × 25 × 25 Å grid, with a spacing of 1 Å. And finally, RosENet was used for predicting the absolute binding affinity (pK_D_), defined as the negative logarithm of the dissociation or inhibition constant, i.e., – log(K_D_) or – log(K_i_), respectively.

**Figure 1.**
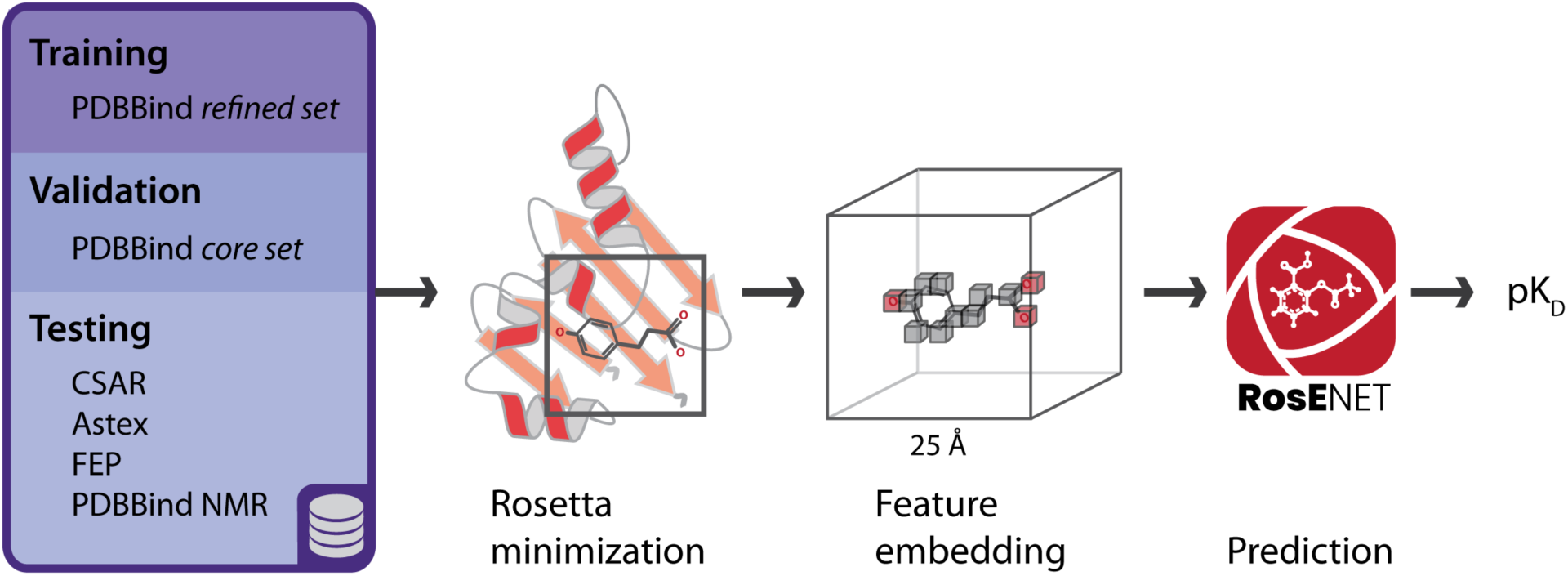
Schematic representation of the RosENet workflow. The PDBBind *refined set, core set* and several other datasets were used, respectively, for training, validating and testing the model. Each protein – ligand complex was first minimized using Rosetta software. The molecular energies and descriptors of the binding pocket were extracted and voxelized (feature embedding). RosENet was used to predict the binding affinity (pK_D_).

### Feature extraction and embedding

One novelty of our approach is the use of molecular energies as a source for 3D features of the complexes. In a 3D set-up, many approaches implicitly use force-fields for ligand docking, where the position of the ligand is optimized by minimizing the energy of a molecular force field. We further exploited this information by directly integrating the energy terms as feature maps. We focused on the Rosetta all-atom force field and considered the attractive, repulsive, electrostatic and implicit solvation energies between pairs of non-bonded atoms (**Figure 2**). The pairwise interactions between the protein and the ligand were clearly visible in their respective voxelized energy maps where high intensity voxels were collocalized (red and blue surfaces, respectively, in **Figure 2b**).

**Figure 2.**
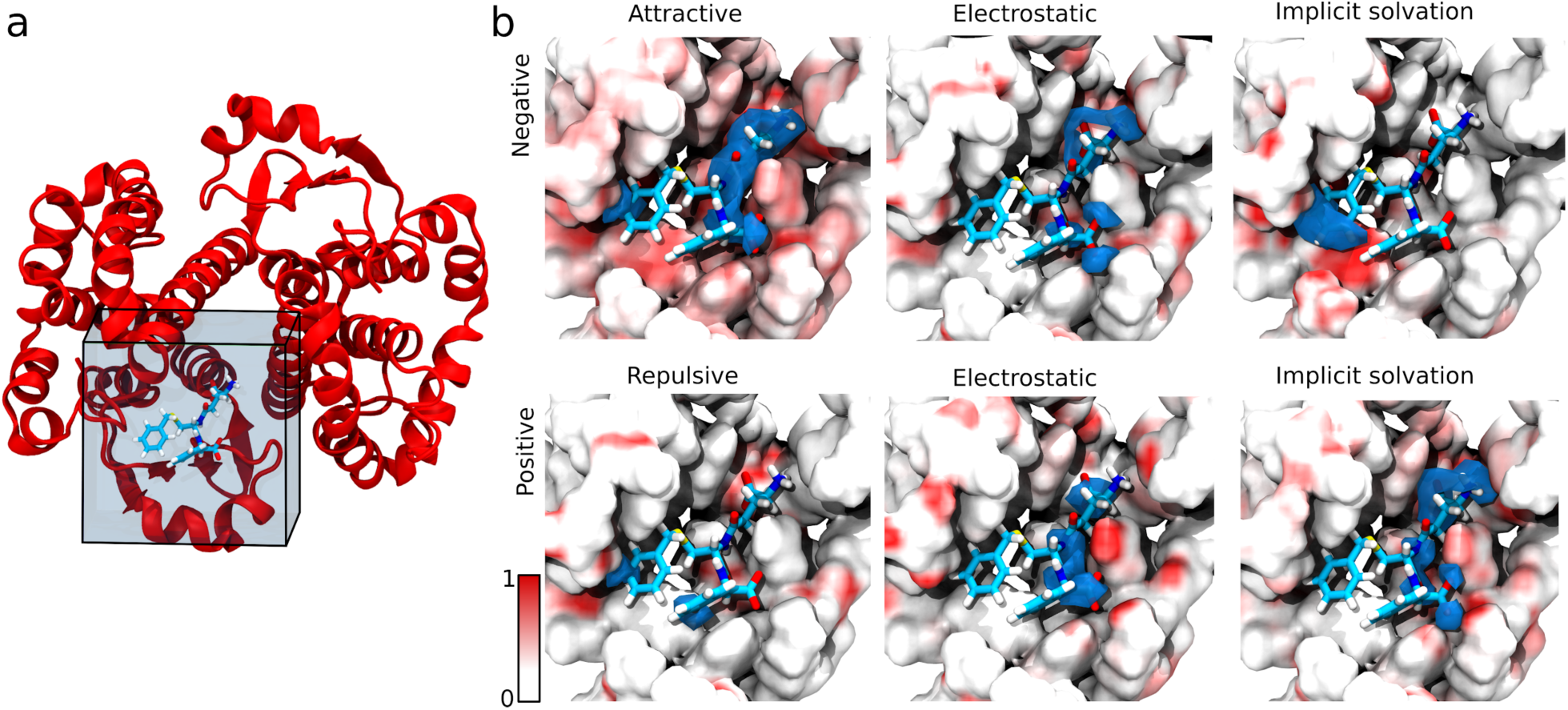
Representative voxelization of the energy features. a. Human Glutathione in complex with the TER117 inhibitor (PDB id: 10GS). The protein and ligand are represented as a red ribbon and light blue sticks, respectively. The cube delineates the binding site used for the 25 × 25 × 25 Å grid, and b. Voxelized representation of energy features. The upper panels display the negative energy terms and the lower panels display the positive energy terms. The protein is shown as a surface representation, where the color scale ranges from 0 (white) to 1 (red); see legend. The ligand is shown with light blue sticks and the 0.65 isocontour of the voxelized ligand is illustrated as a blue surface.

For the molecular descriptors, we followed a similar approach as used for KDeep,^19^ and selected a subset of the following 4 molecular descriptors from AutoDock Vina:^7^ i) aromatic carbon, ii) hydrogen bond acceptor, iii) positive ionizable, and iv) negative ionizable. These descriptors provide the chemical nature of the atomic interactions that we hypothesized would complement the energy terms. For example, aromatic carbons are involved in important and specific interactions, such as π–π stacking.

### Training and validation

RosENet was initially trained with the PDBBind *refined set* minus the *core set* for 300 epochs. We trained twenty replicas from scratch, and for each replica, the model that minimized the validation error of the *core set* was saved. This usually occurred after approximately 270 epochs. The model with the overall least validation error (RMSE of 1.27, 95% Confidence interval (CI): [1.18, 1.38]) was chosen for further analysis. We observed that the predictions tended to be more accurate for complexes with binding affinity in the medium range (RMSE of 0.88 for pK_D_ = 6 to 8). Conversely, the prediction for weak and strong binders decreased in accuracy, with an RMSE of 1.45 and 1.69 for pK_D_ < 6 and pK_D_ > 9, respectively. The *core set* can be further divided into the CASF-2013 subset,^27,28^ which is commonly used for benchmarking computational tools. For this subset, the error was slightly higher (RMSE of 1.53), but the overall correlation coefficient remained the same for both sets (R: 0.8). These results are comparable to alternative Deep Learning approaches.^12,19,20^

Due to the apparent dataset bias, we retrained RosENet with an extended dataset, combining the previous complexes from the PDBBind *refined* set and added complexes from the BindingMOAD^29^ with pK_D_ < 4 or pK_D_ > 9.^29^ This newly generated model achieved an RMSE of 1.26 (95% CI: [1.16, 1.37]) (**Figure 3**). The accuracy was closer to the average for binding affinities on the low range (pK_D_ < 6, RMSE 1.28) and medium range (pK_D_ >= 6 and pK_D_ <= 9, RMSE 1.04). The higher binding affinities (pK_D_ > 9, RMSE 1.84) were still not as well represented as the rest of the range. For the CASF-2013 subset, the error and correlation coefficient remained similar (RMSE 1.55, R: 0.77).

**Figure 3.**
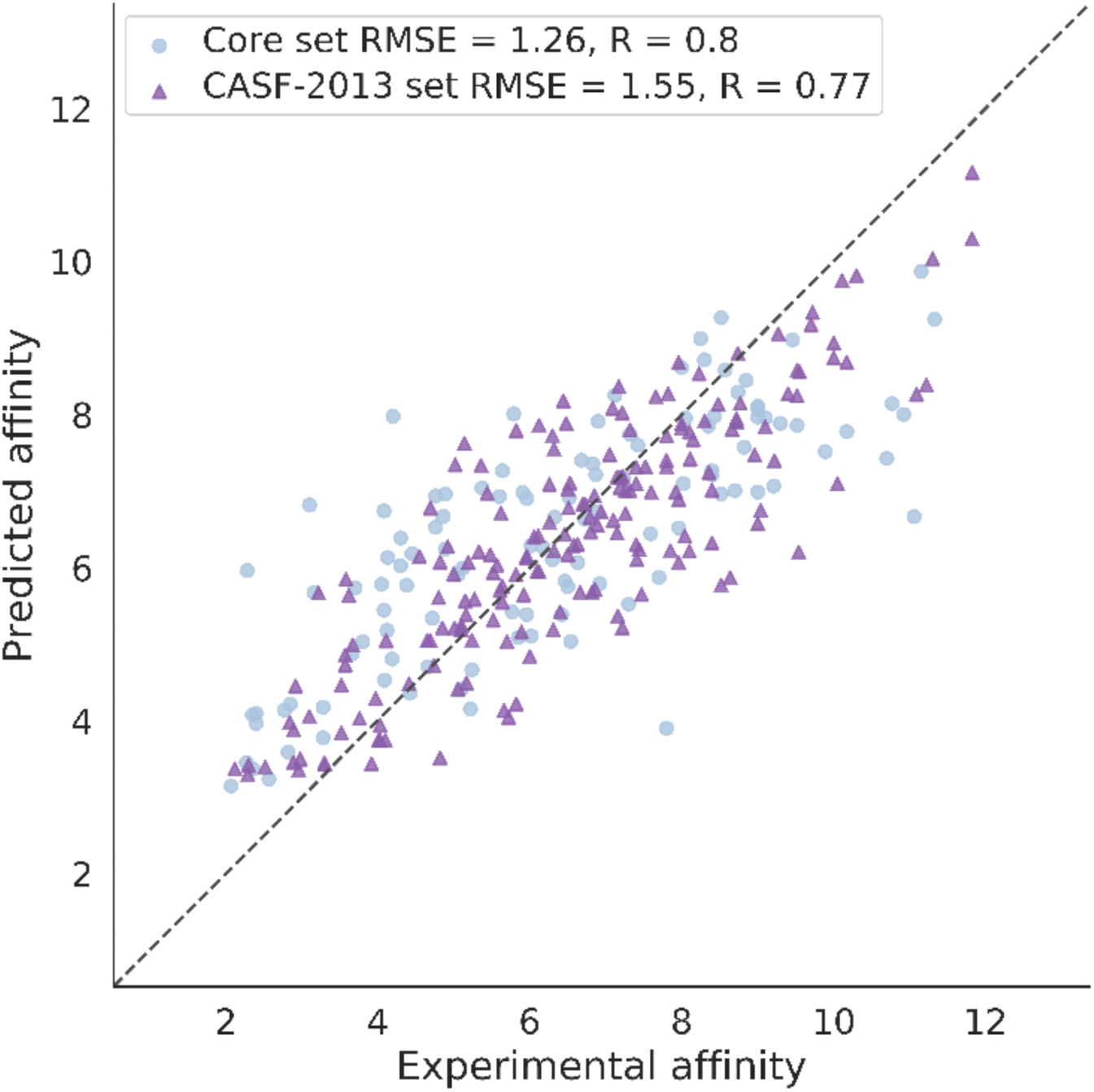
Binding affinity predictions for the validation set. The PDBBind *core set* is shown with light blue circles and the CASF-2013 subset is highlighted with dark purple triangles. The Root Mean Square Error (RMSE) and Spearman’s correlation coefficient (R) are reported for the full *core set* and for the CASF-2013 subset (see legend).

Lastly, the effect of the Rosetta relaxation procedure on the prediction error was tested. The PDBBind *core set* was relaxed using a slightly different protocol that allowed less movement for the ligand. The error and correlation coefficient remained similar (RMSE: 1.3, R: 0.8), thus supporting the robustness of the RosENet.

Data from 60 different protein families/targets were used to build the *core set*. When analyzing the RMSE and correlation coefficient for the disaggregated *core set*, the predictions for the majority of the targets were highly correlated with the experimental affinity (Spearman’s correlation coefficient: average R: 0.74, median R: 0.84, **Figure 4**). Only one target reported a negative correlation for the predictions: O-GlcNAcase (BT4395: 12 structures, average pK_D_: 6.21 +/− 1.04, R: −0.04). We did not observe a direct dependence between the prediction error and correlation.

**Figure 4.**
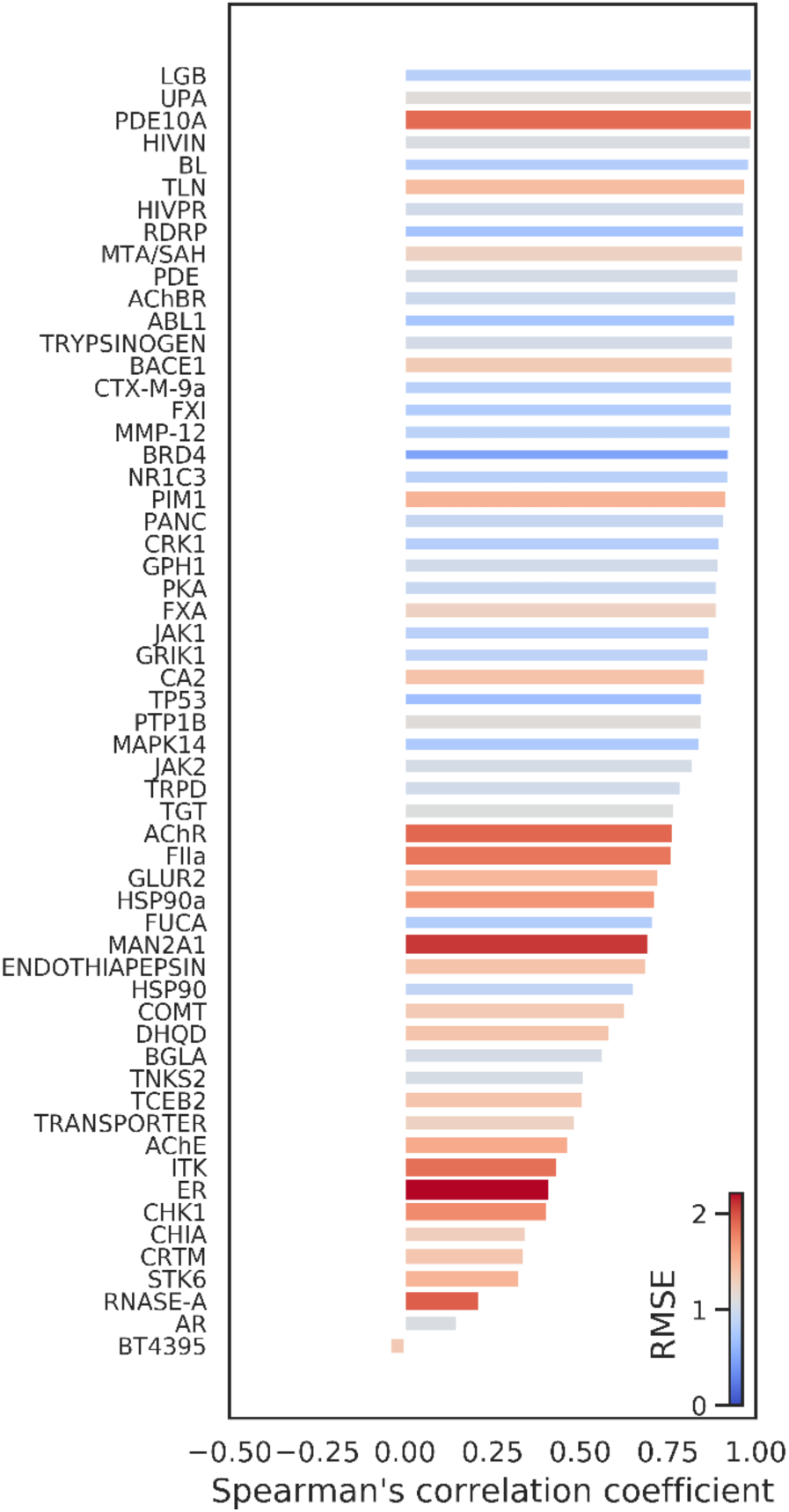
Disaggregated correlation and RMSE for each target of the PDBBind *core set*. The bar length represents the Spearman’s correlation coefficient, and the bar’s width and color, the RMSE within the target cluster.

### Testing

In total, RosENet was tested on 446 structures from 4 different datasets: CSAR, Astex, FEP and PDBBind NMR. These datasets were significantly different from the training data, since only 52 target proteins were similar to a protein present in the PDBBind *refined set* (90% sequence identity). The RMSE and Spearman’s correlation coefficient (R) of the prediction against the true binding affinities were computed for each test set (**Figure 5 and Table S1 in the Supplementary Information (SI) section**). The CSAR dataset was further subdivided into two sets: HQ1 and HQ2.^30,31^ The RMSE increased for HQ1 and HQ2 (1.75, 95% CI: [1.51, 2.21] and 1.43, 95% CI: [1.21, 1.81], respectively). The Astex dataset considerably overlapped with the PDBBind *refined* and *core sets* and therefore was the smallest test dataset with only 17 complexes.

**Figure 5.**
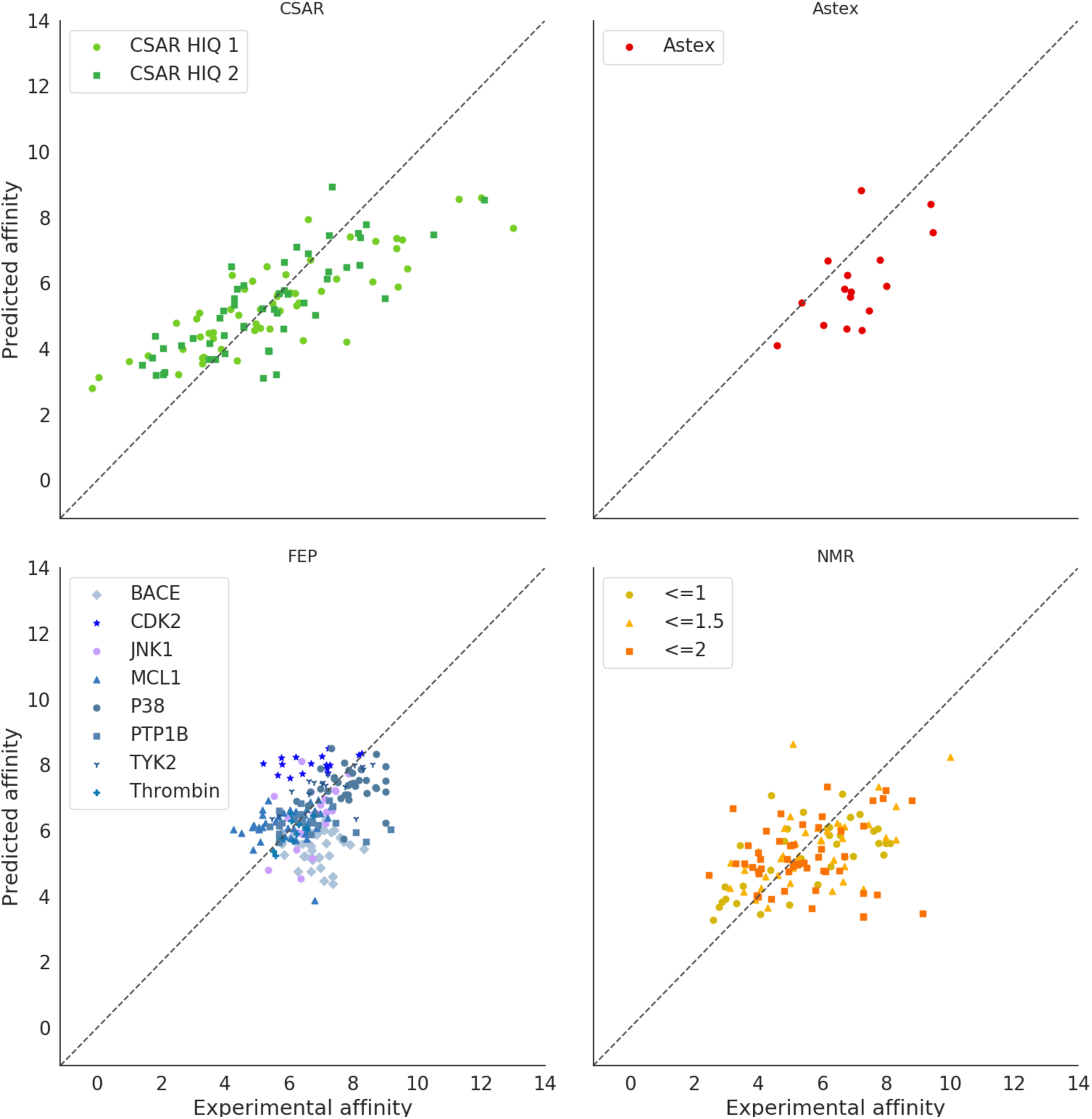
Binding affinity predictions for the test sets. Four datasets, i.e., CSAR, Astex FEP and NMR were used to test RosENet. Subsets are shown with different colors and markers (see legend).

The FEP dataset was composed of eight different proteins for which the binding affinity of 11 to 42 ligands have been measured.^32^ As a whole set, the predictions were good, although it should be noted that the complexes’ binding affinity ranges mainly from 6 to 8, where the predictions seem to be the most accurate. When disaggregating the data, we observed a large variability in the predictions (**Figure S1 in SI**). For example, BACE and P38 achieved very low Spearman’s correlation coefficients (R: 0.03 and R: 0.01 respectively), whereas TYRK2 obtained a much better correlation coefficient (R: 0.64).

We also tested the impact of the experimental structure determination technique and thus extracted structures obtained by Nuclear Magnetic Resonance (NMR) for the PDBBind *general set*. Structures for which the ligand moved by a Root Mean Square Deviation (RMSD) greater than 2 Å after minimization were excluded from the dataset. A 2 Å cut-off is commonly used for docking benchmarks. We observed a considerable decrease of predictive performance in the complexes that were displaced by maximum 2.0 Å after minimization compared to all of those displaced by maximum 1.5 Å. This difference is appreciable in both the RMSE and Spearman’s correlation coefficient (RMSE: 1.51, 95% CI: [1.34, 1.72], R: 0.42 and RMSE: 1.35, 95% CI: [1.15, 1.58], R: 0.59, respectively). No difference was measured when comparing thresholds of 1 Å and 1.5 Å, respectively.

Lastly, we compared RosENet to two recent methods that are also based on CNNs and for which the codes were available: OnionNet and Pafnucy. Since OnionNet was trained on the entire PDBBind *general set*, we could only consider structures from CSAR and FEP that did not overlap with it. RosENet and Pafnucy achieved the best performance for the CSAR dataset (RMSE: 1.62, 95% CI: [1.34, 2.00], R: 0.70) and FEP dataset (RMSE: 0.93, 95% CI: [0.84, 1.03], R: 0.39), respectively (**Figure 6**). OnionNet did not perform well on the FEP dataset (RMSE: 1.53, R: −0.02).

**Figure 6.**
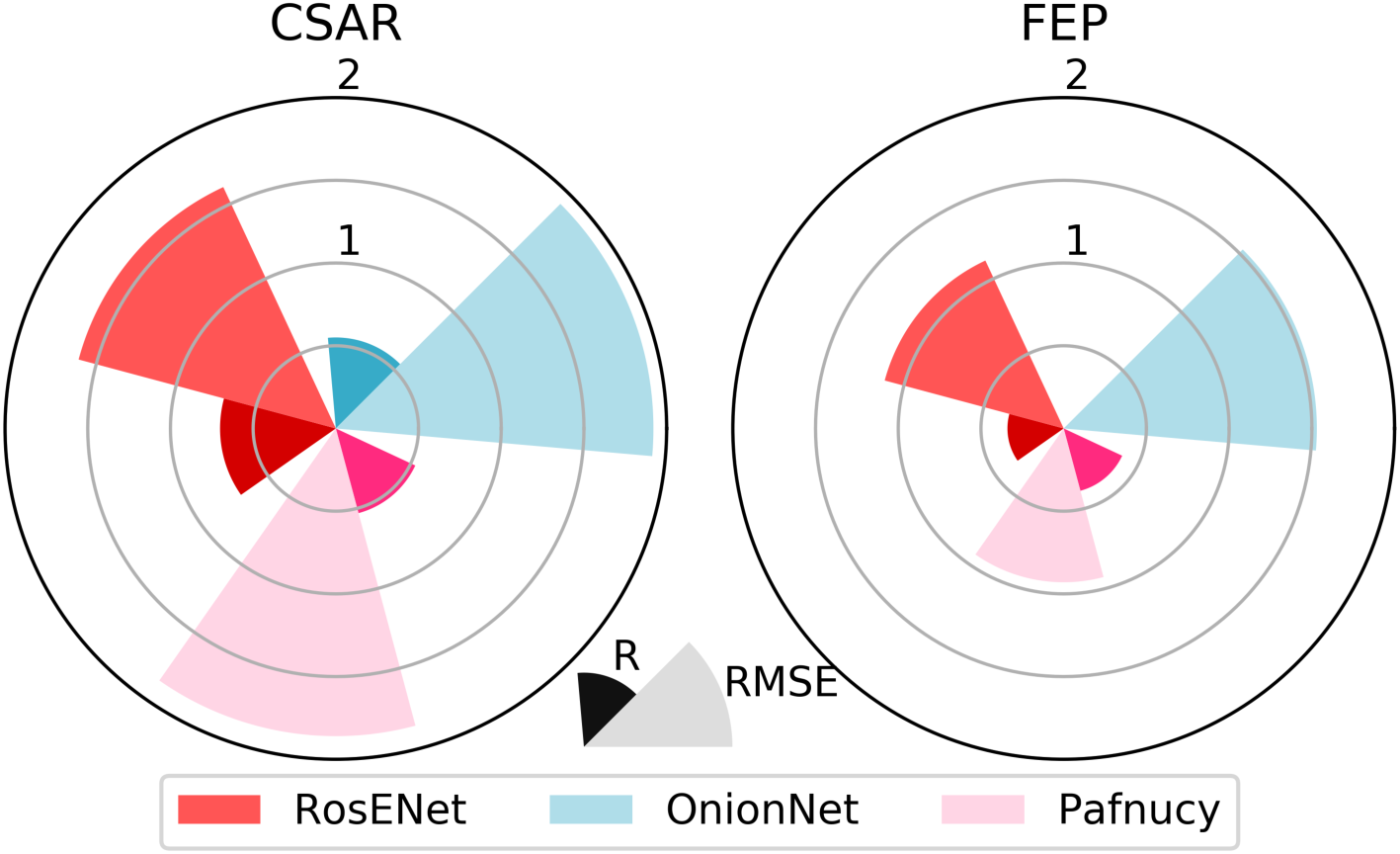
Circular bar plot comparing RosENet to OnionNet and Pafnucy for prediction accuracy. The RMSE and Spearman’s correlation coefficient (R) are reported in red, cyan and pink, respectively, for RosENet, OnionNet and Pafnucy. The RMSE is shown with the lighter shades of color, and R with the darker shades of color. Note that the correlation coefficient for OnionNet on the FEP was negative and therefore omitted on the plot (R: −0.02, p-value: 0.69)

### Robustness testing

It has been shown that the accuracy of some Machine Learning models can be very sensitive to small changes in the input data. To test the robustness of RosENet, OnionNet and Pafnucy, we carried virtual screening experiments. Data generated from these experiments would also be more representative of a computational drug discovery pipeline. Each complex in the PDBBind *core set* and test set (CSAR and Astex) was docked 20 times. The poses with lowest Rosetta Energy score and smallest RMSD were considered for building the two docking test datasets. Overall, RosENet achieved the best performance (RMSE: 1.62, R: 0.70) (**Figure 7**). The prediction error for the PDBBind *core set* increased when selecting the lowest energy and lowest RMSD structures for RosENet (RMSE: 1.26 95% CI: [1.16, 1.37] and RMSE: 1.42, 95% CI: [1.30, 1.54], respectively) and OnionNet (RMSE: 1.32., 95% CI: [1.22, 1.44] and RMSE: 1.52, 95% CI: [1.41, 1.66], respectively). However, the error for Pafnucy remained comparable for both datasets (RMSE: 1.67, 95% CI: [1.54, 1.82] and RMSE: 1.62, 95% CI: [1.50, 1.77], respectively). For the Test set, the prediction error remained comparable for all models independent of the docking dataset.

**Figure 7.**
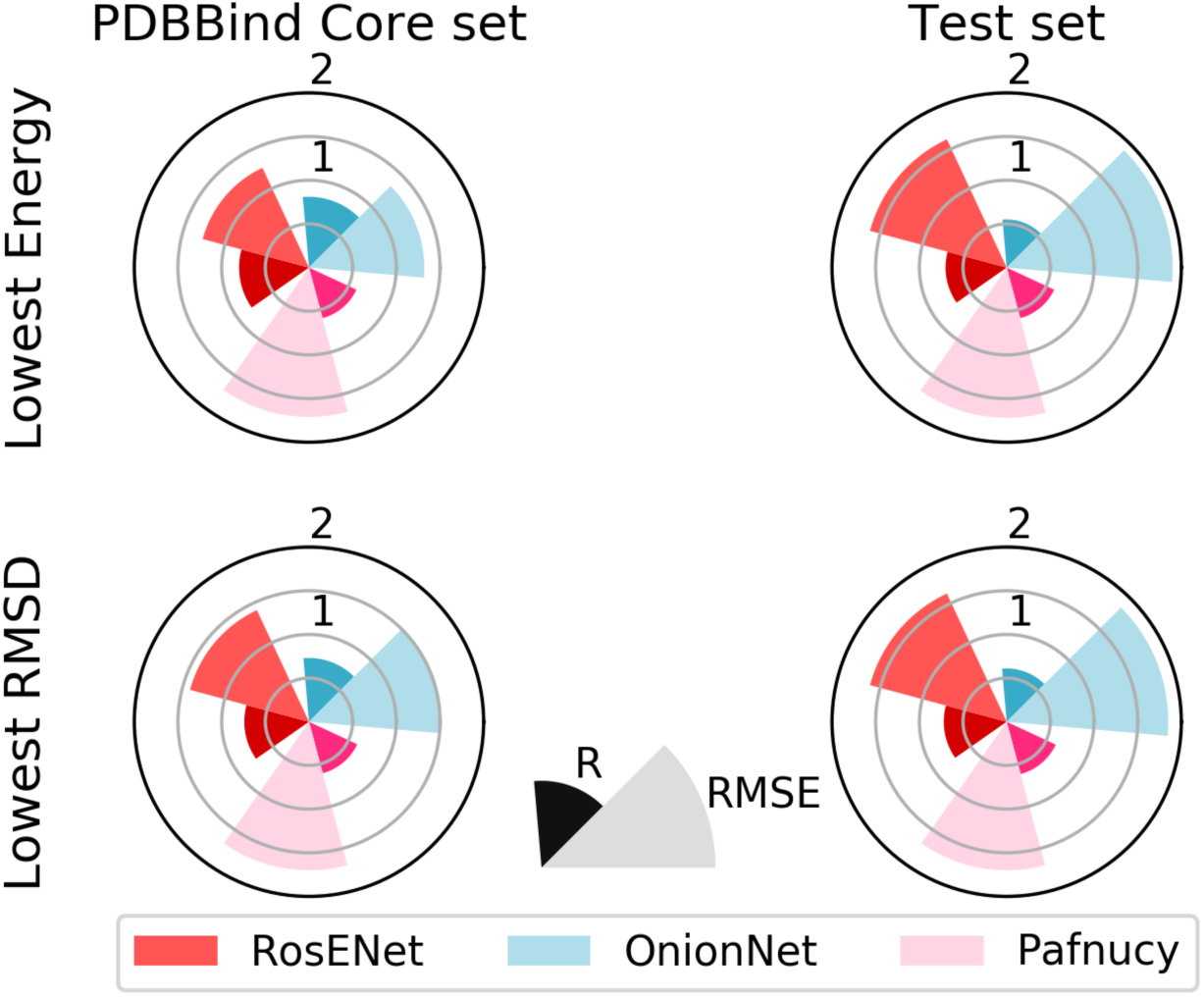
Circular bar plot comparing RosENet to OnionNet and Pafnucy for the virtual screen experiments. The RMSE and Spearman’s correlation coefficient (R) are reported in red, cyan and pink, respectively, for RosENet, OnionNet and Pafuncy. The RMSE is shown with the lighter shades of color, and R with the darker shades of color.

We lastly compared the correlations between the predictions of RosENet and OnionNet or Pafnucy. The predictions from RosENet correlated slightly better with the ones of Pafnucy (R: 0.65) than with OnionNet (R: 0.44), but remained low in both cases (**Figure 8**). These noticeable differences would suggest that ensemble methods could improve the accuracy of the predictions. We estimated the theoretical minimum RMSE on the combined test sets, by choosing the most accurate prediction from each network. Pafnucy predominantly made the predictions that were closest to the experimental measure (105 closest predictions), followed by RosENet (90) and OnionNet (65). By combining all three networks, an RMSE of 0.80 and R of 0.84 (p-value: 1.88e-66) could theoretically be achieved.

**Figure 8.**
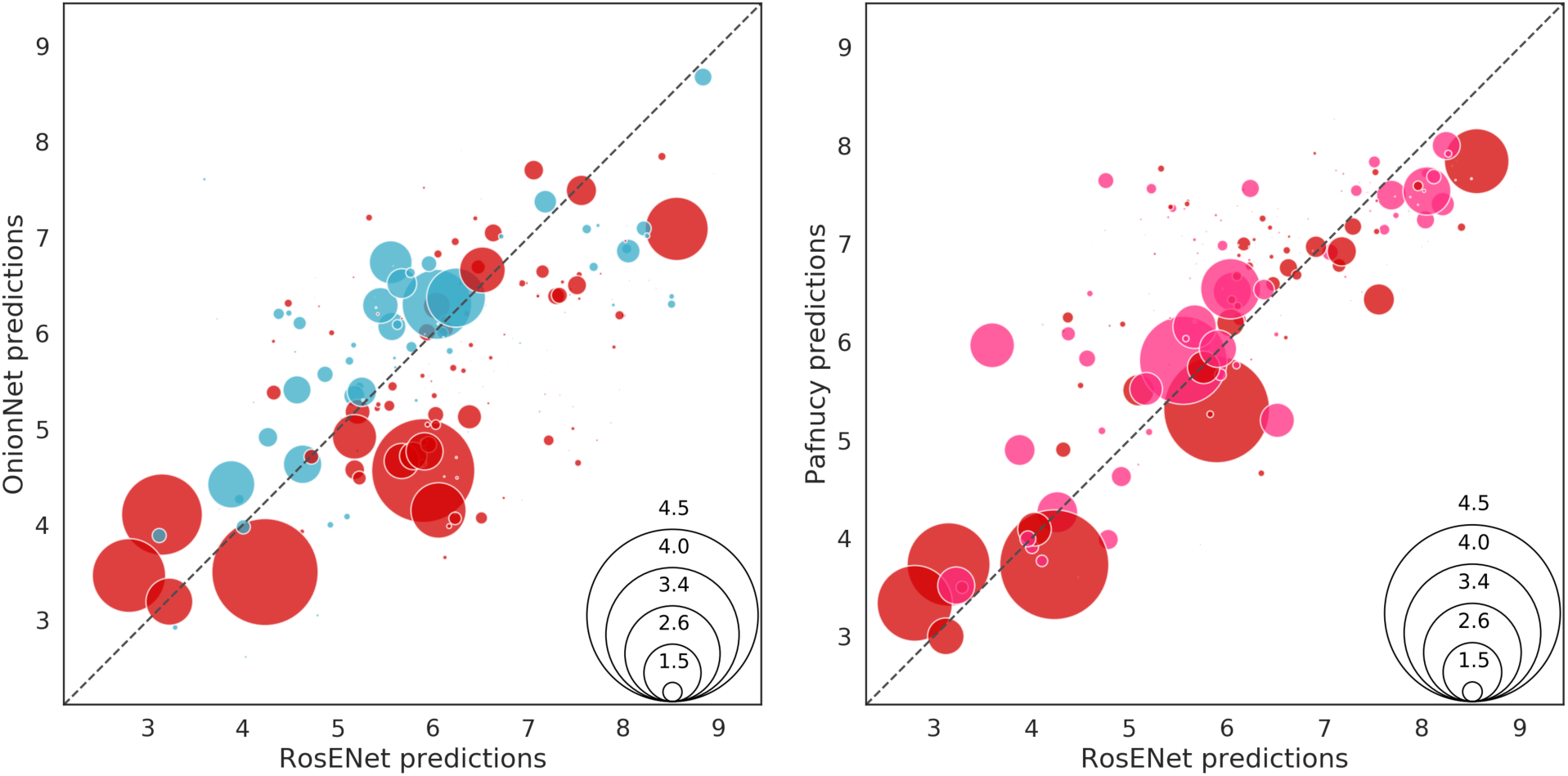
Predictions for RosENet compared to Pafnucy and OnionNet. Each circle represents the prediction for a complex. The radius of the circle is scaled to the RMSE of the best prediction and the color indicates which network achieved the best prediction, i.e., red: RosENet, cyan: OnionNet, and pink: Pafnucy.

### Feature ablation and the effect of Network architecture

We tested the effect of feature ablation and network architecture on the quality of the predictions. A feature ablation was first carried out comparing the performance of the models using only: i) the energy terms, ii) the 4 molecular descriptors (4 HTMD), or iii) the 8 molecular descriptors (8 HTMD) provided by HTMD.^33^ A smaller network architecture (SqueezeNet)^34^ was then tested, that was similar to the one used for KDeep.^19^

Overall, RosENet showed the best performance. The model using only 4 molecular descriptors (4 HTMD) achieved a comparable level of accuracy, whereas the use of additional molecular descriptors (8 HTMD) marginally affected the accuracy (**Figure 9**, top panel and **Table S3 in SI**). The greatest decrease in performance was observed when using the energy terms alone. The 8 HTMD features performed better with SqueezeNet architecture than with ResNet (**Figure 9**, lower panel, and **Table S4 in SI**). For the SqueezeNet architecture, the addition of energy features systematically had a deleterious effect on the performances. We performed a T-test to evaluate the statistical significance of differences in the predictions between RosENet and the other configurations. For this analysis, all the test datasets were merged and a total of 7 T-tests were carried out. In all cases, RosENet achieved a significantly lower Mean Square Error (MSE), with the highest p-value being approximately 5% (**Table S5 in SI**).

**Figure 9.**
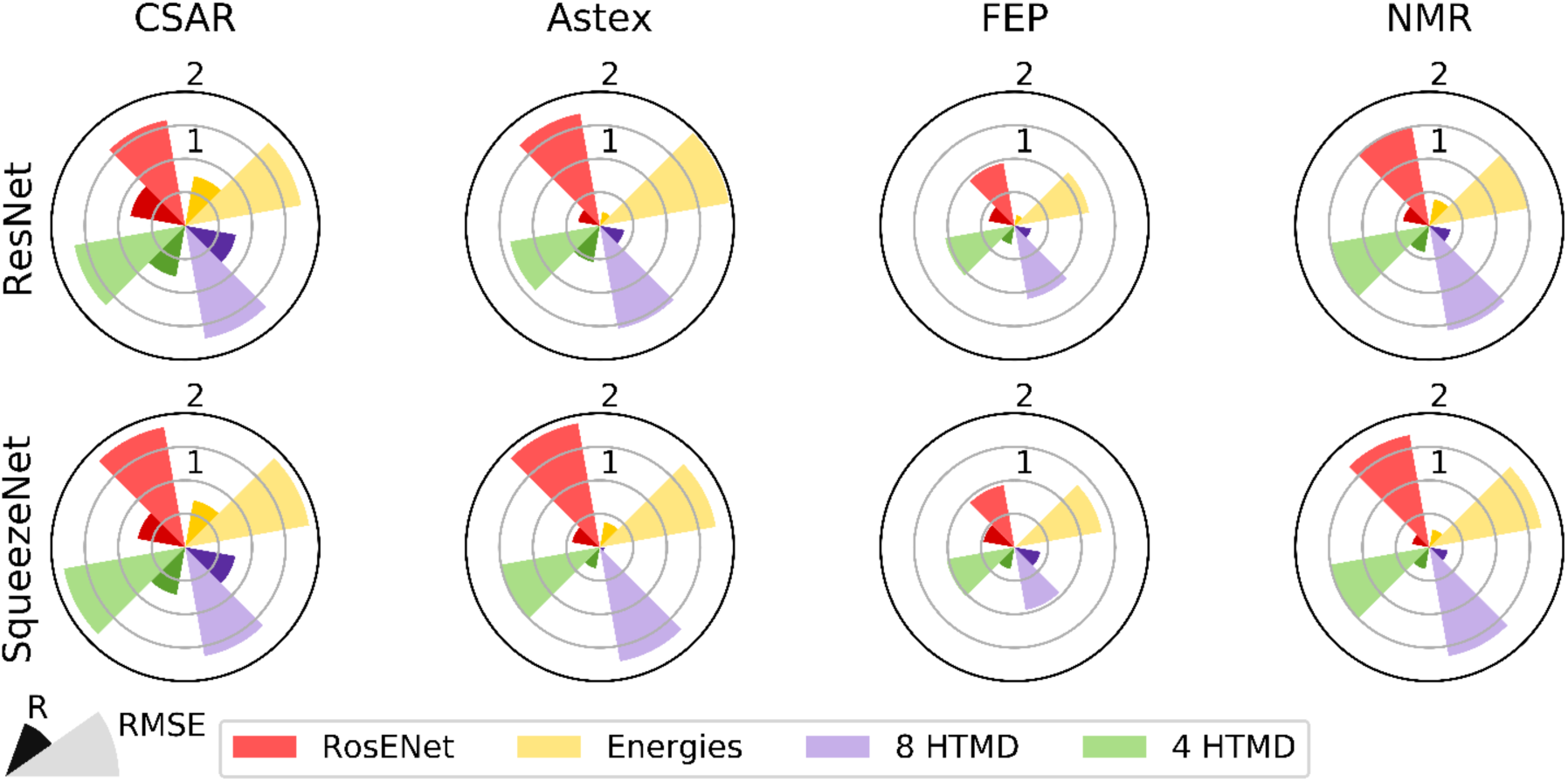
Circular bar plot comparing the effect of feature ablation and network architecture. Each set of features is represented with a different color (see legend). The RMSE is shown with the lighter shades of color and Spearman’s correlation coefficient (R) with the darker shades of color. The ResNet and SqueezeNet architectures were used for the models in the upper and lower panels, respectively.

## DISCUSSION

In this study, we developed RosENet, a 3D Convolutional Neural Network that combines voxelized molecular mechanics energies and molecular descriptors in order to improve the prediction of the binding affinity of protein – ligand complexes. We have shown that the molecular energies can be embedded into a 3D grid and complement molecular descriptors when training a CNN. When presented unseen data, RosENet was able to better generalize than previously proposed methods (**Figures 6 and 7**). We hypothesized that the energy terms provide critical information for estimating the binding affinity (e.g., the solvation energy) that cannot be captured using only molecular descriptors. Models using the SqueezeNet architecture and trained with the energy terms did not perform as well, indicating that the relation between the energy features and the experimental binding affinity cannot be easily represented in a shallower network. Furthermore, the residual connections that appear in the ResNet architecture might also help identify important differences in energies within the complexes instead of simply the absolute energies.

The parameters of the ligand were determined using the procedure provided by Rosetta, which has the advantage of being fast. This parameterization remains coarse, especially for the atomic charges. A more refined procedure, such as those used for molecular dynamics, might help to further improve the prediction of the binding affinity. One drawback, however, would be that this would be computationally more intensive. In this study, only a subset of energies computed by Rosetta was considered. In the future, other energy terms can be added, but this would also significantly increase the complexity and size of the feature maps. To deal with that, a feature selection procedure would probably first be required. It should be noted that our voxelization procedure could also be extended to energy terms computed with other force fields (e.g., CHARMM,^35^ AMBER.^36^ OPLS^37^ or GROMOS^38^) or docking packages (e.g., AutoDock Vina^7^).

During validation and testing, we observed that the performance of the prediction was strongly dependent on the range of the binding affinity and the protein family. Such a behavior has been reported for most of the previously proposed models. Typically, a broader range of sampled binding affinity values leads to an increase in the RMSE of the predictions, due to a degradation of the prediction for high and low binding affinities. As described in previous studies,^19^ the accuracy of the predictions also depends on the protein family. This type of behavior is typically indicative of overfitting. It was also discussed that the PDBBind dataset is strongly unbalanced.^39,40^ Some protein families, such as the HIV integrase with 282 structures, are overrepresented in the PDBBind *refined set* (**Figure S2 in SI**). Furthermore, the distribution in pK_D_ is not uniform (**Figure S3 in SI**), but follows a quasi-normal distribution centered around 6.4 with a standard deviation of 2. This could explain why our model performed better for predicting binding affinities in the median range; in particular, since the RMSE was used as a loss function during training and would thus maximize the likelihood of an assumed normal distribution. By integrating structures from the BindingMOAD in an effort to rebalance the PDBBind training set, we observed a small improvement for the lower values of pK_D_. Balancing or resampling procedures could thus be applied in the future to prioritize underrepresented structures. When comparing RosENet to OnionNet and Pafnucy, we observed that certain models performed better for some complexes. One could take advantage of this by training and combining several models using ensemble methods (e.g., boosting).

## CONCLUSIONS

We demonstrated that molecular energies can be voxelized and combined with molecular descriptors in order to predict the binding affinity of protein – ligand complexes. In the future, this framework can be extended to other features extracted from biophysical models, e.g., molecular dynamics simulations. The dynamics of the protein and ligand are known to have a significant effect on the binding affinity. In order to further improve the development of Deep Learning approaches, an important effort would be required for building well-balanced training, validation and testing datasets. Working in that direction, we have already collected and curated almost 400 structures that can be used for testing models. We have also built a new dataset of structures solved by NMR, where the ligand RMSD after minimization was used to assess the quality of the structures. Lastly, we have shown the potential for improving the predictions by using ensemble methods.

## MATERIALS AND METHODS

This section is divided into six parts. In the first part, we will present the different datasets used for training, validating and testing our models. The features computed for the protein – ligand complex, the voxelization and the representation procedure used for embedding will then be described. The architectures and training procedure of the different convolutional neural networks will follow. And finally, we will present the analyses that were used to compare the accuracy of the different models.

### A. Datasets

#### A.1 Training and validation set

##### PDBBind

The PDBBind dataset was used for training and validating our models. PDBBind is divided into three concentric sets. The *general set* contains the experimental data for the 3D structure and the experimentally measured binding affinity of 19 588 biomolecule complexes. From the *general set*, a curated *refined set* of 4 463 protein – ligand complexes of high structural resolution (< 2.5 Å and an R-factor higher than 0.25) with accurate binding measurements (1 pM < K_d_ / K_i_ < 10 uM) was extracted. Finally, a 90% sequence similarity cut-off was used to group the structures in the *refined set* into 58 different clusters. Complexes from 58 representative clusters were chosen and constituted the *core set*. The binding affinity pK_D_ was defined as the negative logarithm of the dissociation or inhibition constants, -log(K_d_) or -log(K_i_), respectively.

The models were trained with the *refined set* (combining versions 2016 and 2018), excluding the *core set* structures. The *core set* was used as the validation set.

#### A.2 Test datasets

In total, the four datasets described below were used for testing our models. Structures already present in the training or validation sets were excluded. The full list of PDB IDs for each dataset is provided in the Supporting Information section.

##### A.2.1 Community Structure-Activity Resource (CSAR)

The Community Structure-Activity Resource (CSAR) was designed for testing and improving computational methods for drug discovery. In this study, the CSAR NRC-HiQ datasets was chosen. This dataset was part of the CSAR 2010 exercise and was composed of two sets: structures discovered between 2004 and 2006, and structures from 2007 and 2008. Both sets were balanced in terms of different statistical properties of the structures. We excluded the structures that were already used in other datasets (for training, validation or testing), which resulted in 55 structures for CSAR HIQ (Set 1) and 49 structures for CSAR HIQ (Set 2).

##### A2.2 FEP

The FEP dataset was used for a Free Energy Perturbation study^32^ and was composed of the following eight proteins: BACE, CDK2, JNK1, MCL1, P38, PTP1B, Thrombin and Tyk2. In addition, 199 ligands were included, thus forming 40 complexes with BACE, 16 with CDK2, 21 with JNK1, 42 with MCL1, 34 with P38, 23 with PTP1B, 11 with Thrombin and 12 with Tyk2.

##### A.2.3 PDBBind 2018 NMR structures

Complexes obtained by Nuclear Magnetic Resonance (NMR) from the PDBBind 2018 dataset were extracted. A filtered version of the NMR dataset was produced by restricting it to structures with a buried Solvent-Accessible Surface Area (SASA) ratio larger than 15%, a molecular mass of maximum 1000 g/mol, and a RMSD after minimization smaller or equal to 2 Å. This yielded 126 structures.

##### A2.4 Astex Diverse dataset

The Astex Diverse dataset is a manually curated dataset composed of 85 complexes with drug-like ligands. After excluding structures overlapping with previous datasets, 17 complexes were kept.

#### A.3 Features

Two different types of features were considered in this study, namely molecular descriptors and molecular energies. Molecular descriptors were defined using AutoDock Vina atom types. Molecular energies were computed with the Rosetta full-atom force field.

##### A.3.1 Molecular descriptors

Eight different molecular descriptors were generated with the Python library HTMD^33^ (Table 1). From these eight descriptors, the following four were selected to be combined with the molecular energies: i) aromatic carbon, ii) hydrogen bond acceptors, iii) positive ionizable, and iv) negative ionizable. Each atom was assigned a binary value for every descriptor.

**Table 1.**
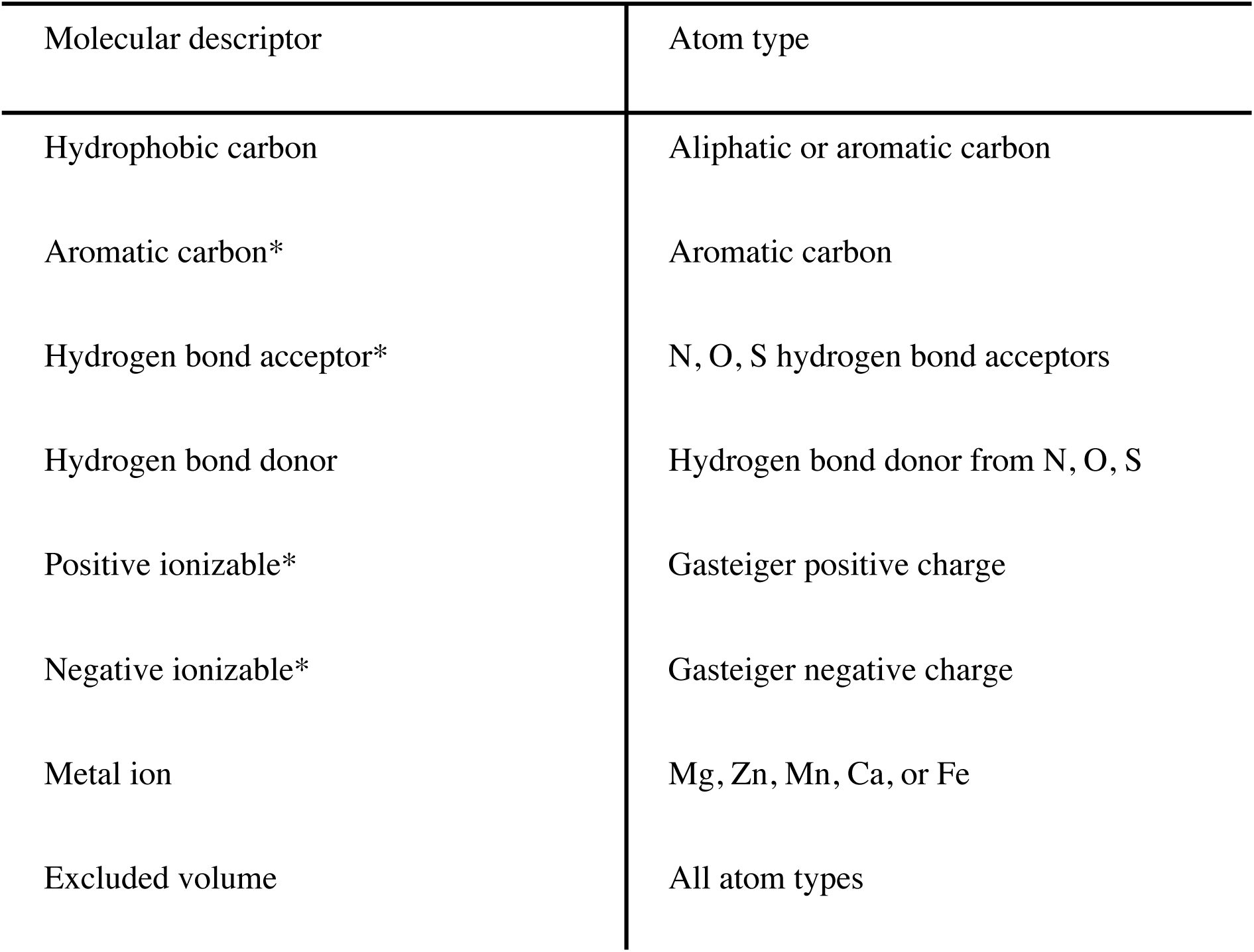
Molecular descriptors computed with HTMD. The descriptors marked with an asterix were combined with the molecular energies.

##### A.3.2 Molecular energies

The ligands were parameterized using Rosetta’s *molfile_to_params*.*py* script. The interface between the ligand and protein was then minimized using RosettaScript. The XML description of the full procedure is available on the github repository. In total 10 complexes were generated for each protein – ligand complex, and the full-atom energies for the structure with the lowest energy score were extracted. The four main full-atom energy terms in REF15 were considered: attractive (fa_atr), repulsive (fa_rep), electrostatic (fa_elec) and implicit solvation (fa_sol).

Emulating the separation of positive and negative charges in the molecular descriptors, we split the energies into 6 terms: attractive, repulsive, positive electrostatic, negative electrostatic, positive implicit solvation, and negative implicit solvation. Each energy term was then normalized to the interval of 0 to 1 over the entire dataset, in order to match the range of the rest of molecular descriptor features.

##### A3.3. Representation

A 3D grid representation was used for embedding the features of the protein – ligand complex. The features for the protein and ligand were voxelized separately. For each feature, a 25 × 25 × 25 Å cubic grid with a spacing of 1 Å and centered around the geometric center of the ligand was created. The position of each atom within the grid was mapped to a voxel and the value of the corresponding feature was assigned to the voxel.

The values of each voxel were then spatially distributed using the function:

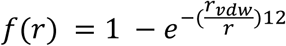

where r is the distance between the grid point and the atom, and r_vdw_ is the Van der Waals radius of the atom. Only the largest contribution was considered for voxels where the values from several atoms overlapped. These grids were combined to form a 25 × 25 × 25 image with one channel for each feature. Three sets of images were created with the following channels for the protein and ligand: 8 molecular descriptors, 4 molecular descriptors, and 4 molecular descriptors with 6 molecular energies.

#### A.4 Architectures of the Convolutional Neural Networks

Two different Convolutional Neural Networks were evaluated, namely ResNet and SqueezeNet.

##### A.4.1 ResNet

To predict the binding affinities of the complexes, we used a version of the well-established neural network architecture ResNet^25^ (**Figure 10** and **Table 2**).

**Table 2.**
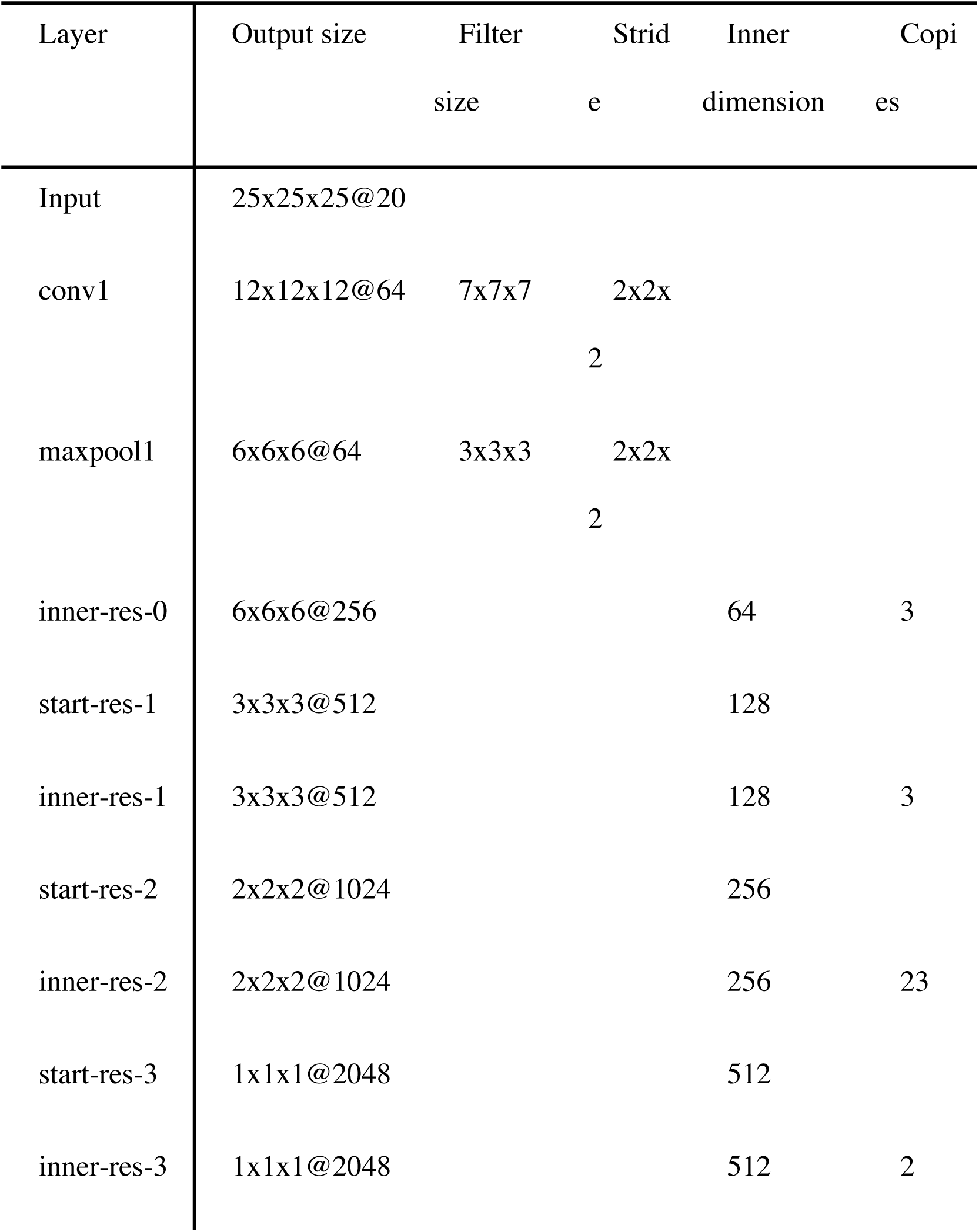
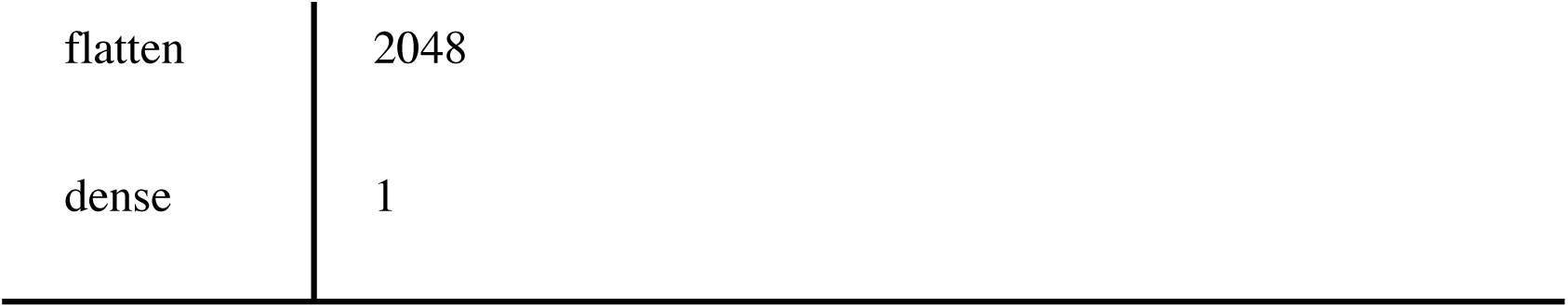
Architecture of the ResNet 101-layer one.

| Layer | Output size | Filter size | Stride | Inner dimension | Copies |
| --- | --- | --- | --- | --- | --- |
| Input | 25x25x25@20 |  |  |  |  |
| conv1 | 12x12x12@64 | 7x7x7 | 2x2x2 |  |  |
| maxpool1 | 6x6x6@64 | 3x3x3 | 2x2x2 |  |  |
| inner-res-0 | 6x6x6@256 |  |  | 64 | 3 |
| start-res-1 | 3x3x3@512 |  |  | 128 |  |
| inner-res-1 | 3x3x3@512 |  |  | 128 | 3 |
| start-res-2 | 2x2x2@1024 |  |  | 256 |  |
| inner-res-2 | 2x2x2@1024 |  |  | 256 | 23 |
| start-res-3 | 1x1x1@2048 |  |  | 512 |  |
| inner-res-3 | 1x1x1@2048 |  |  | 512 | 2 |

|  |  |
| --- | --- |
| flatten | 2048 |
| dense | 1 |

**Figure 10.**
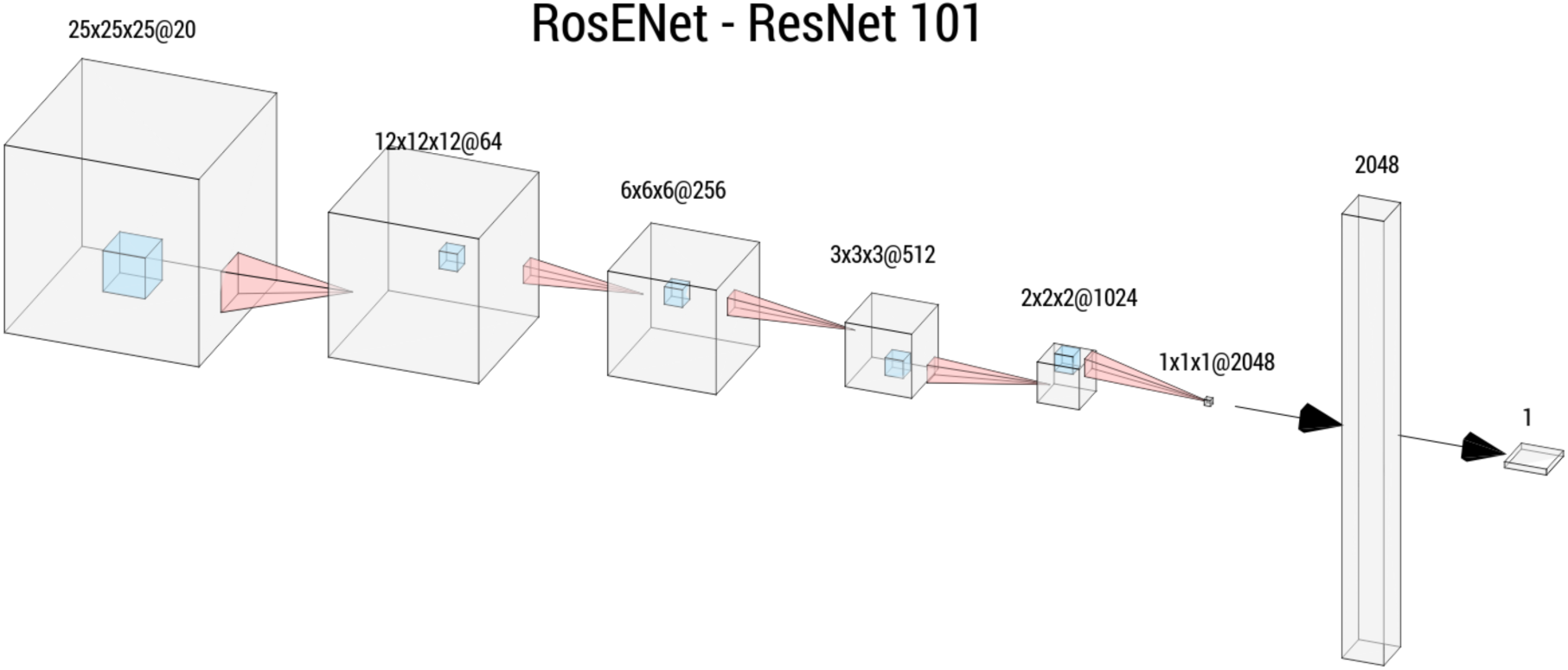
Schematic representation of the architecture of RosENet. The input was composed of a 25 × 25 × 25 image with 20 channels. The blue cubes show the convolution filter size.

ResNet is composed of “residual modules.” Each module has three layers, with filter sizes 1×1×1, 3×3×3 and 1×1×1, respectively. Residual modules include skip connections, with the assumption that the modules should not learn all attributes of the data, but only the perturbations between input and output. Residual modules do not reduce the size of the images, so ResNet assembles them in blocks separated by max pooling layers that perform the downsampling. There are also skip connections between contiguous blocks, skipping the separating max pooling layers.

In our architecture, the first convolutional layer was followed by a ReLU, whereas the rest of the convolutional layers used in the residual modules, were preceded by a ReLU, as recommended in.^41^

##### A.4.2 SqueezeNet

The SqueezeNet architecture is composed of fire modules, which are two-layered modules of convolutional layers (**Figure 11** and **Table 3**). The first layer of the module is the squeeze layer, a convolutional layer with a filter size 1 × 1 × 1 and a small number of filters. This squeeze layer is supposed to constrain the network to a more compact representation in order to find the features that are more important and reject the noisier ones. The second layer is the expand layer, composed of two side-by-side convolutional layers, one with a size 1 × 1 × 1 and the other one with a size 3 × 3 × 3, with a larger channel count than the squeeze layer. This layer is dedicated to performing transformations over the “squeezed” representation. These side-by-side convolutions are then concatenated to form an output with a much larger channel count. Each fire module maintains the dimensions of the images, and either maintains or increases the number of channels.

**Table 3.**
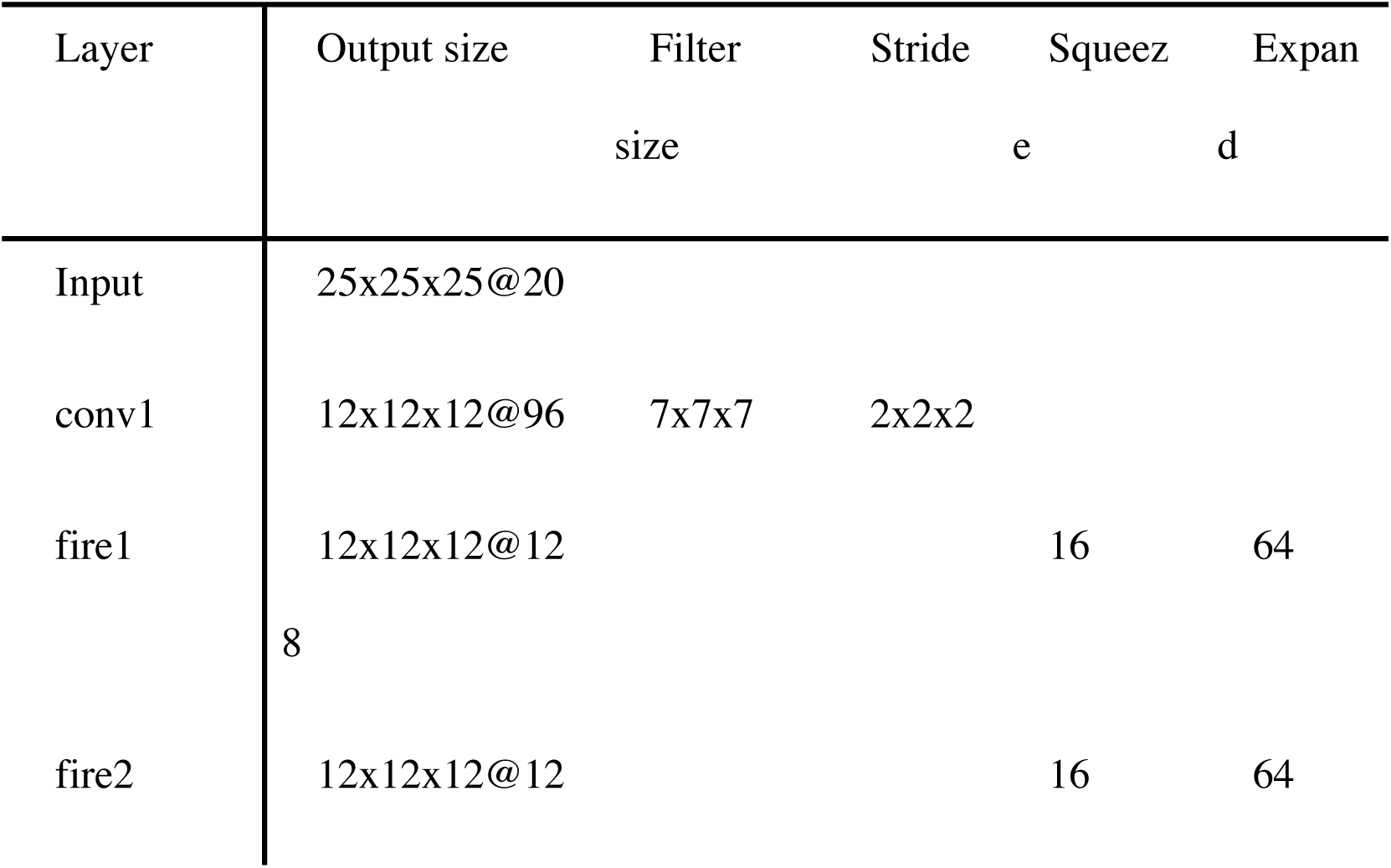
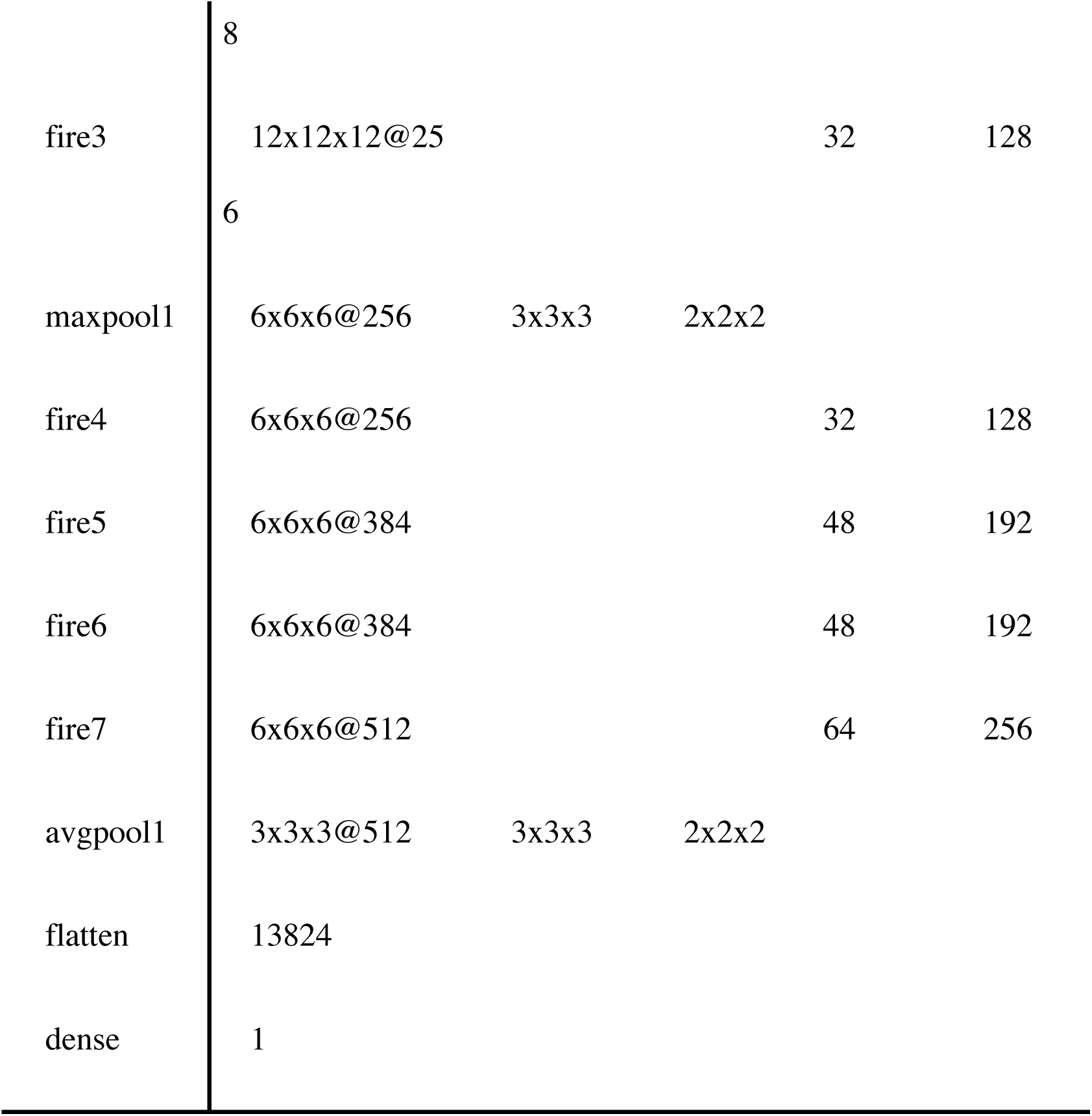
Architecture of the SqueezeNet.

**Figure 11.**
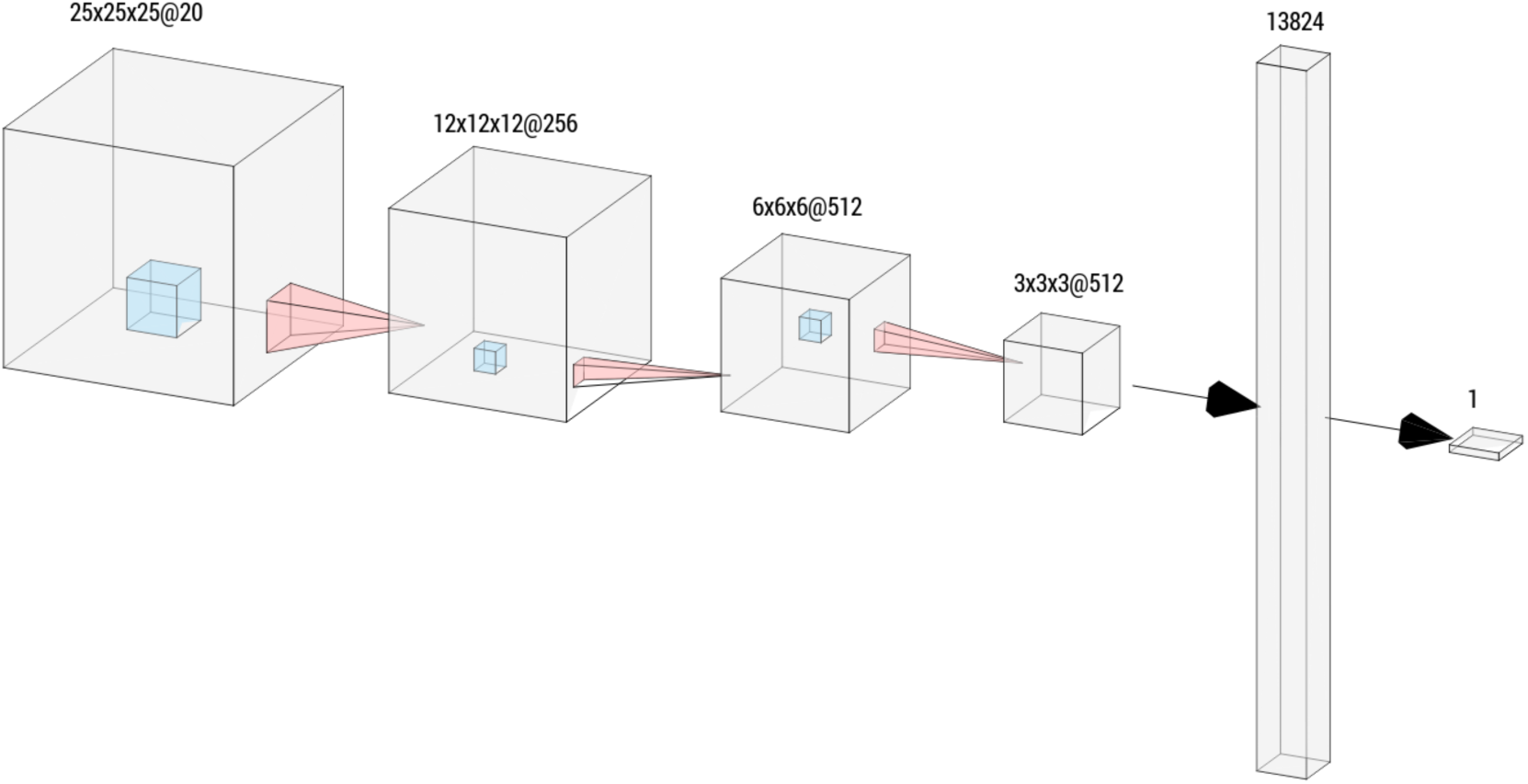
Schematic representation of the architecture of SqueezeNet. The input was composed of a 25 × 25 × 25 image with 20 channels. The blue cubes show the convolution filter size.

The implementation of these neural networks is available on our GitHub site. https://github.com/DS3Lab/RosENet/tree/master/models

#### A.5 Training

The loss function was defined as the Root Mean Square Error (RMSE):

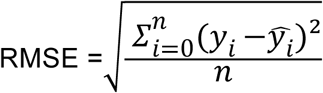

where *ŷ*_*l*_ is the predicted binding affinity and *y*_*i*_ is the true binding affinity.

Batch Normalization (BN) and Dropout were not used, since they significantly worsened the performance of our models for all of our experiments.

For training, the AdaM optimizer^42^ with its default parameters and a learning rate of 10^4^ was used. We trained the networks for 300 epochs using a batch size of 128. The variables are initialized with the default settings of Tensorflow, which corresponds to the Glorot uniform initializer.^43^ In order to guarantee the rotation invariance of our models, each image was augmented by rotating it by 90°, providing 24 additional data points per protein – ligand complex.

#### A.6 Analyses

In addition to the RMSE and Spearman’s correlation, the standard deviation of the linear regression was calculated in order to quantify the quality of the predictions:

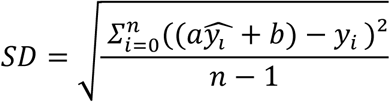

where *ŷ*_*l*_ is the predicted binding affinity, *y*_*i*_ is the true binding affinity, and *a* and *b* are the slope and interception of the linear regression line of the *ŷ*_*l*_ and *y*_*i*_.

## ASSOCIATED CONTENT

Rosenet_SI.pdf: Supporting figures and tables

complexes.zip: list of PDB id and binding affinities for all datasets

## AUTHOR INFORMATION

### Author Contributions

**The manuscript was written with the contributions of all authors. All authors have given their approval of the final version of the manuscript**.

### Funding Sources

**TL would like to gratefully acknowledge the support of the Swiss National Science Foundation (grant P3P3PA_174356)**.

### Notes

## ABBREVIATIONS

CNN: convolutional neural network
RMSE: root mean square error
R: Spearman’s correlation coefficient
NMR: nuclear magnetic resonance
K_I_: inhibition constant
K_D_: dissociation constant
IC_50_: half maximal inhibitory concentration
pK_D_: binding affinities.

## SYNOPSIS

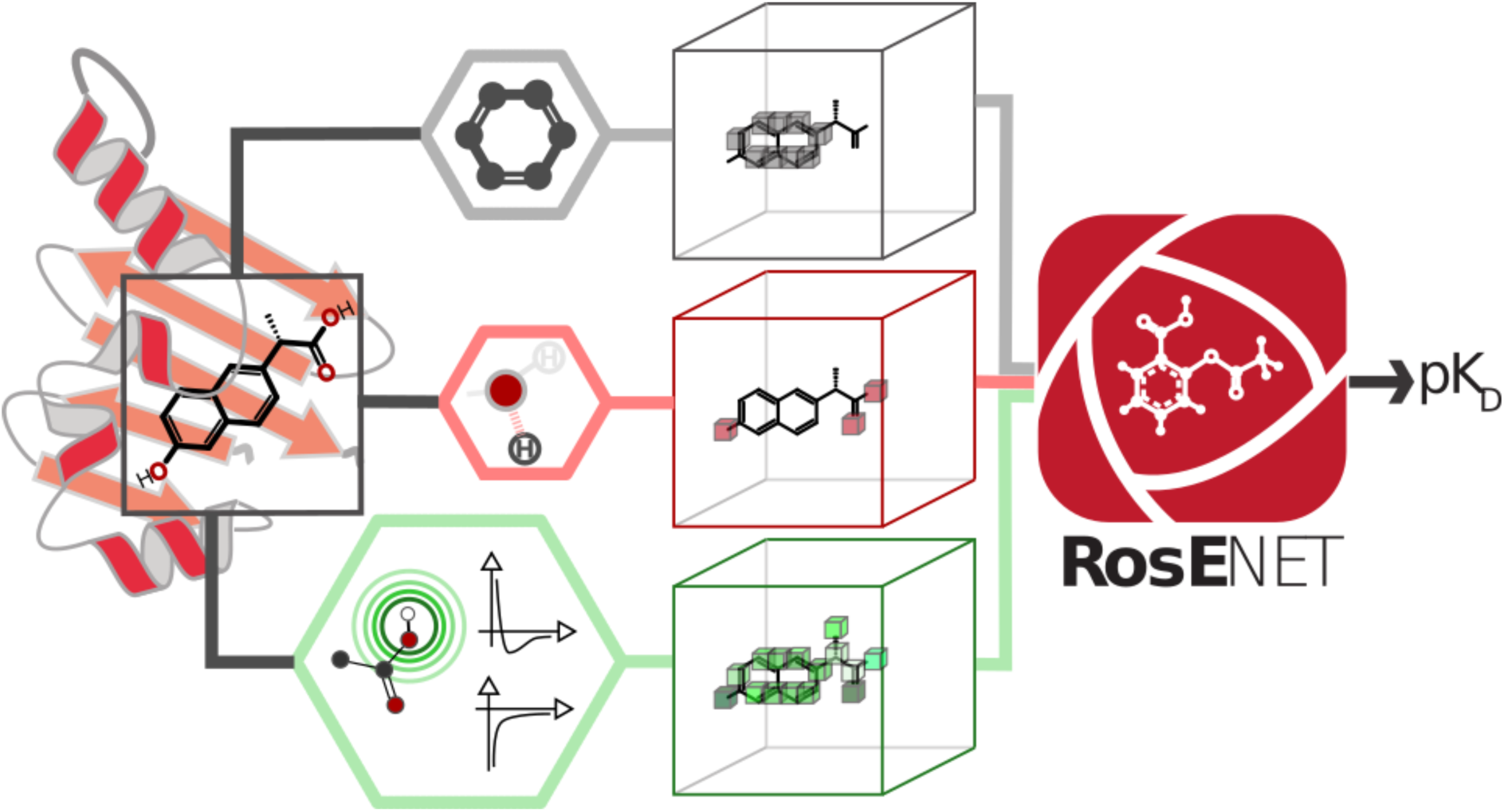

## Notes

https://github.com/DS3Lab/RosENet

### Competing Interest Statement

The authors have declared no competing interest.

