## Supporting information for "RosENet: Improving binding affinity prediction by leveraging molecular mechanics energies with a 3D Convolutional Neural Network"

**SUPPLEMENTARY INFORMATION**

Hussein Hassan-Harrirou[1], Ce Zhang[1], Thomas Lemmin[1,2,*]

[1]DS3Lab, System Group, Department of Computer Sciences, ETH Zurich, CH-8092 Zurich, Switzerland
[2]Institute of Medical Virology, University of Zurich (UZH), CH-8057 Zurich, Switzerland

### Table of contents

**Figure S1. Disaggregated binding affinity predictions for FEP dataset.**

**Figure S2. Heatmap of binding affinity $pK_D$ for the 80 largest clusters.**

**Figure S3. Distribution of binding affinities $pK_D$ in PDBBind dataset.**

**Table S1. Disaggregated RMSE and Spearman's correlation correlations for all test sets.**

**Table S2. Binding affinity predictions for RosENet, OnionNet and Pafnucy.**

**Table S3. Effect of feature ablation on the RMSE and Spearman's correlation coefficient (R) of the binding affinity predictions.**

**Table S4. RMSE and Spearman's correlation coefficient (R) of the binding affinity predictions for all tests when using SqueezeNet.**

**Table S5. T-test comparing the performance of RosENet to models trained during the feature ablation and network architecture analysis.**

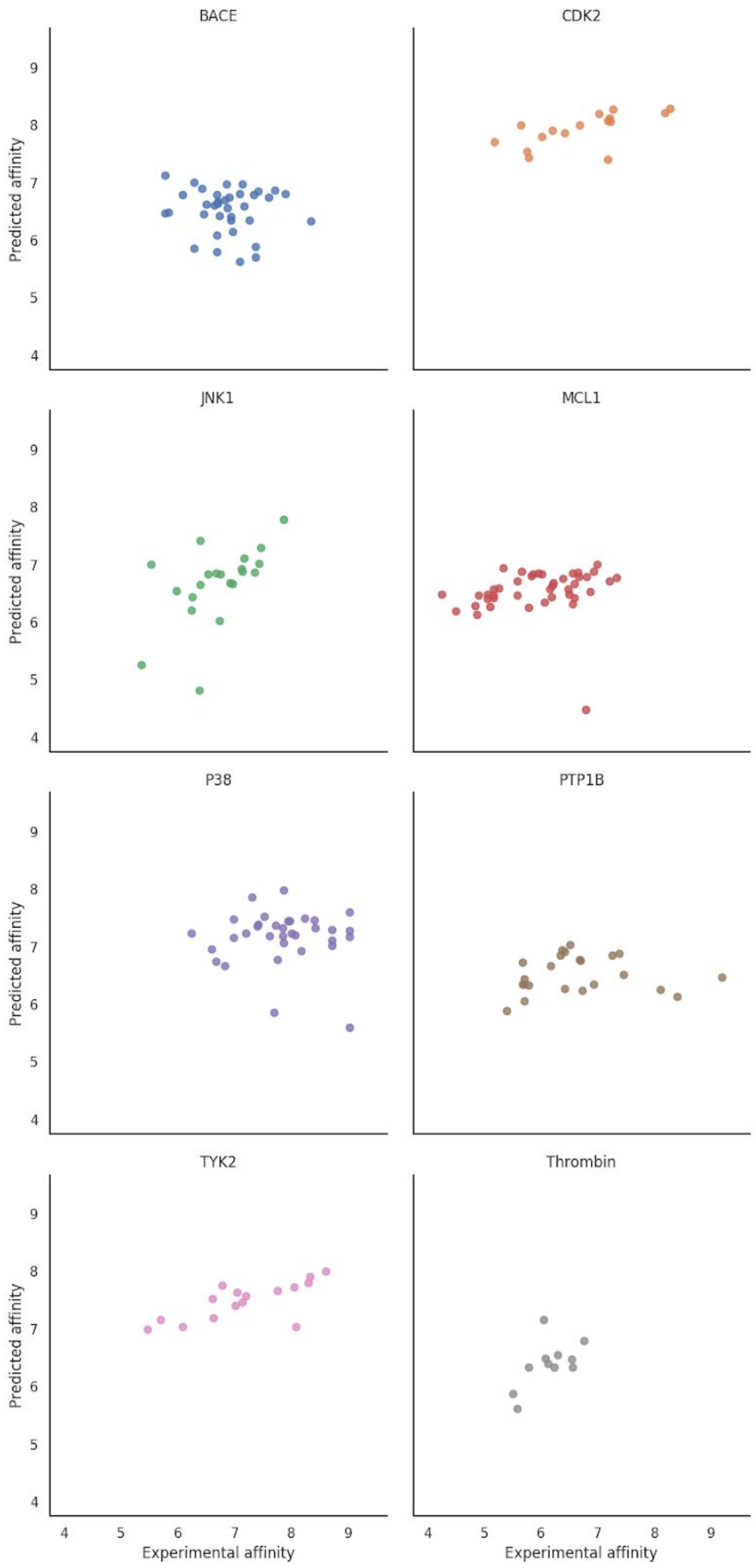


**Figure S1. Disaggregated binding affinity predictions of the FEP dataset.** The target name is indicated in the title.

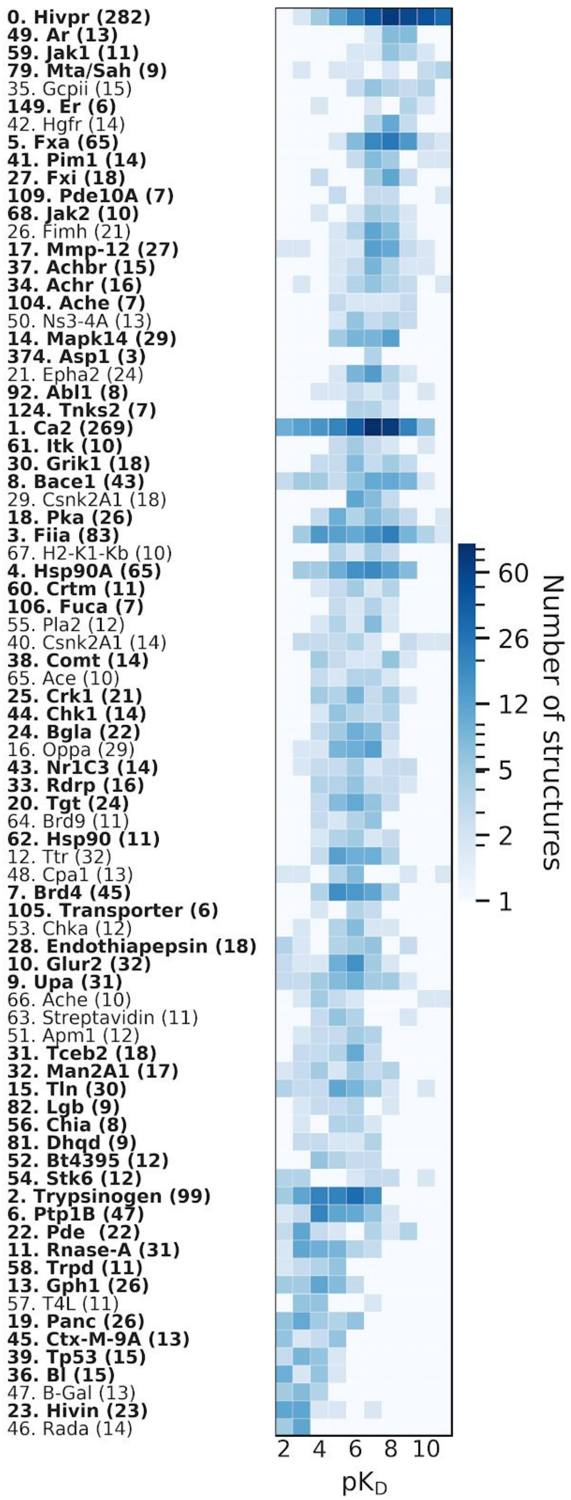


**Figure S2. Heatmap of binding affinity pK$_D$ for the 80 largest clusters.** The clusters are sorted by average pK. The first number indicates the ranking of the cluster sizeThe colors indicate the number of structures present in each interval of pK$_D$ (logarithmic scale). The clusters from which structures were chosen for the PDBBind *core set* are highlighted in bold.

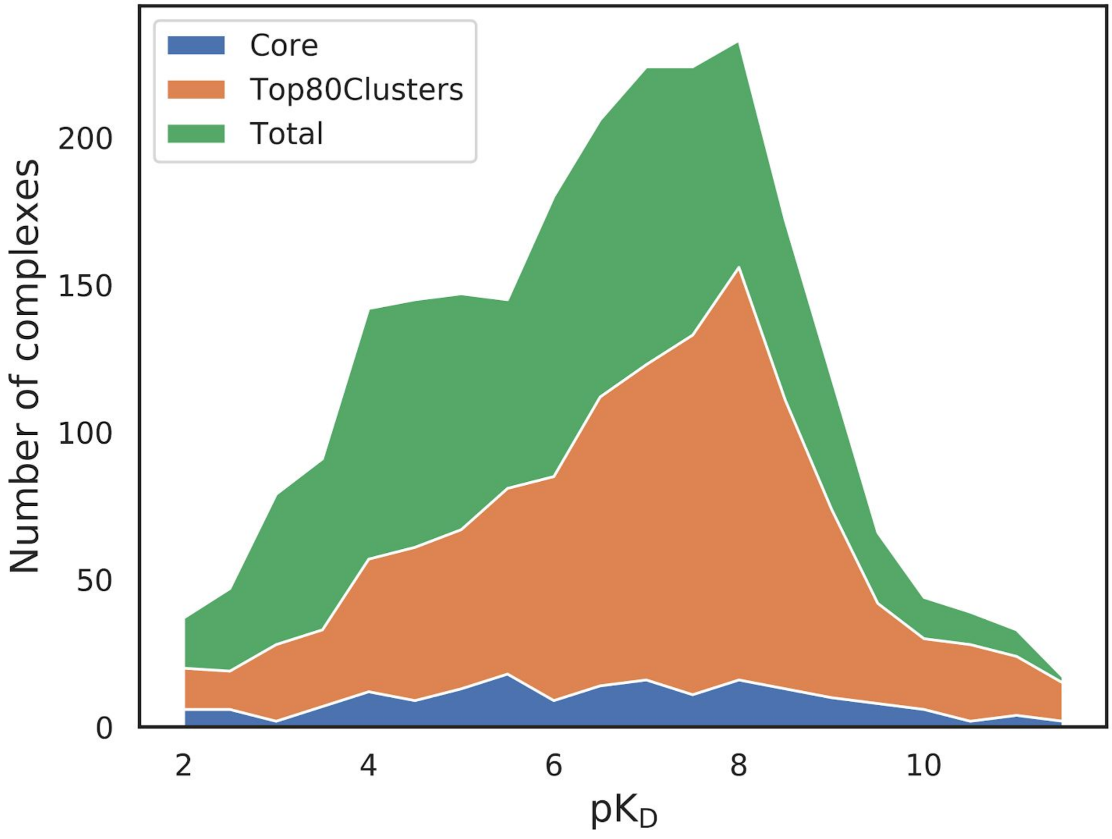


**Figure S3. Distribution of binding affinities $pK_D$.** The PDBBind *core set* is shown in blue, the 80 largest clusters in orange and the *refined set* in green.

**Table S1. Disaggregated RMSE and Spearman's correlation coefficient for all test sets.**

| NMR | | | | | FEP | | | | |
|---|---|---|---|---|---|---|---|---|---|
| | RMSE | R | p-val | SD | | RMSE | R | p-val | SD |
| <=1 Å | 1.45 | 0.46 | 6.00E-03 | 0.97 | BACE | 0.79 | -0.07 | 7.00E-01 | 0.15 |
| <=1.5 Å | 1.41 | 0.51 | 4.00E-06 | 0.86 | CDK2 | 1.43 | 0.72 | 2.00E-03 | 0.05 |
| <=2 Å | 1.5 | 0.41 | 2.00E-06 | 0.87 | JNK1 | 0.59 | 0.65 | 2.00E-03 | 0.31 |
| CSAR | | | | | MCL1 | 1.02 | 0.43 | 5.00E-03 | 0.15 |
| | RMSE | R | p-val | SD | P38 | 1.1 | 0.05 | 8.00E-01 | 0.21 |
| HIQ 1 | 1.75 | 0.85 | 3.00E-16 | 0.69 | PTP1B | 0.98 | 0.11 | 6.00E-01 | 0.1 |
| HIQ 2 | 1.43 | 0.83 | 3.00E-13 | 0.92 | Thrombin | 0.44 | 0.42 | 2.00E-01 | 0.11 |
| Astex | | | | | TYK2 | 0.8 | 0.71 | 2.00E-03 | 0.05 |
| | RMSE | R | p-val | SD | | | | | |
| | 1.7 | 0.34 | 1.00E-01 | 1.63 | | | | | |

RMSE: Root Mean Square Error
R: Spearman's correlation coefficient
p-value: Significance of measured correlation
SD: Standard Deviation of the Linear Regression

**Table S2. Binding affinity predictions for RosENet, OnionNet and Pafnucy for virtual screening experiments.** Results for structures with the lowest energy (minimal energy) and smallest RMSD to ground truth (minimal deviation) are reported. The best performances are highlighted in bold.

| | Datasets | RosENet | | | | OnionNet | | | | Pafnucy | | | |
|---|---|---|---|---|---|---|---|---|---|---|---|---|---|
| | | RMSE | R | p-val | SD | RMSE | R | p-val | SD | RMSE | R | p-val | SD |
| Minimal energy | PDBBind Core | **1.26** | 0.80 | 5e-67 | 0.81 | 1.32 | **0.81** | 1e-70 | 0.78 | 1.67 | 0.65 | 1e-33 | 0.64 |
| | CSAR | | | | | | | | | | | | |
| | HIQ 1 | **1.58** | **0.68** | 4e-6 | 0.45 | 2.10 | 0.42 | 1-e2 | 0.97 | 1.87 | 0.54 | 5e-4 | 1.05 |
| | HIQ 2 | 1.67 | 0.72 | 5e-3 | 1.81 | **1.31** | **0.93** | 3e-6 | 0.99 | 1.82 | 0.63 | 0.02 | 1.31 |
| | Astex | 1.68 | 0.46 | 0.13 | 1.24 | 1.80 | **0.53** | 7e-2 | 1.54 | **1.40** | 0.11 | 0.72 | 0.35 |
| Minimal RMSD | PDBBind Core | **1.41** | **0.74** | 4e-50 | 1.00 | 1.52 | 0.73 | 2e-49 | 2.07 | 1.62 | 0.65 | 1e-35 | 0.65 |
| | CSAR | | | | | | | | | | | | |
| | HIQ 1 | **1.59** | **0.73** | 2e-6 | 0.42 | 1.95 | 0.58 | 2e-4 | 0.90 | 1.81 | 0.59 | 1e-4 | 0.73 |
| | HIQ 2 | 1.65 | 0.77 | 2e-3 | 1.67 | **1.31** | **0.91** | 1e-5 | 0.70 | 1.73 | 0.60 | 0.02 | 1.20 |
| | Astex | 1.66 | 0.41 | 0.17 | 1.22 | 2.07 | **0.45** | 0.13 | 1.13 | **1.26** | 0.24 | 0.44 | 0.36 |

RMSE: Root Mean Square Error
R: Spearman’s correlation coefficient
p-value: significance of measured correlation
SD: Standard Deviation of the Linear Regression

**Table S2. Effect of feature ablation on the RMSE and Spearman's correlation coefficient (R) of the binding affinity predictions for all tests.** The best performances are highlighted in bold.

| Datasets | RosENet | | | | Energies | | | | 4 HTMD | | | | 8 HTMD | | | |
|---|---|---|---|---|---|---|---|---|---|---|---|---|---|---|---|---|
| | RMSE | R | p-val | SD | RMSE | R | p-val | SD | RMSE | R | p-val | SD | RMSE | R | p-val | SD |
| Astex | 1.7 | 0.34 | 1.00E-01 | 1.63 | 1.96 | 0.21 | 4.00E-01 | 2.28 | 1.36 | 0.55 | 1.00E-02 | 1.27 | 1.56 | 0.37 | 1.00E-01 | 1.32 |
| CSAR | | | | | | | | | | | | | | | | |
| HIQ 1 | 1.75 | 0.85 | 3.00E-16 | 0.69 | 1.89 | 0.8 | 1.00E-13 | 0.65 | 1.79 | 0.84 | 2.00E-15 | 0.84 | 1.89 | 0.82 | 1.00E-14 | 0.85 |
| HIQ 2 | 1.43 | 0.83 | 3.00E-13 | 0.92 | 1.62 | 0.71 | 1.00E-08 | 0.94 | 1.57 | 0.7 | 2.00E-08 | 1.32 | 1.51 | 0.73 | 2.00E-09 | 1.22 |
| NMR | | | | | | | | | | | | | | | | |
| <=1 Å | 1.45 | 0.46 | 6.00E-03 | 0.97 | 1.5 | 0.46 | 6.00E-03 | 0.85 | 1.34 | 0.6 | 2.00E-04 | 0.97 | 1.35 | 0.62 | 8.00E-05 | 1.1 |
| <=1.5 Å | 1.41 | 0.51 | 4.00E-06 | 0.86 | 1.47 | 0.46 | 4.00E-05 | 0.86 | 1.42 | 0.49 | 8.00E-06 | 0.98 | 1.44 | 0.53 | 1.00E-06 | 1.13 |
| <=2 Å | 1.5 | 0.41 | 2.00E-06 | 0.87 | 1.5 | 0.4 | 3.00E-06 | 0.92 | 1.48 | 0.41 | 2.00E-06 | 0.97 | 1.6 | 0.32 | 2.00E-04 | 1.23 |
| FEP | 0.96 | 0.4 | 5.00E-09 | 0.34 | 1.13 | 0.17 | 2.00E-02 | 0.55 | 1.06 | 0.28 | 5.00E-05 | 0.55 | 1.11 | 0.25 | 3.00E-04 | 0.65 |

RMSE: Root Mean Square Error
R: Spearman's correlation coefficient
p-value: Significance of measured correlation
SD: Standard Deviation of the Linear Regression

**Table S3. RMSE and Spearman's correlation coefficient (R) of the binding affinity predictions for all tests when using SqueezeNet.** The best performances are highlighted in bold.

| Datasets | Energies + 4HTMD | | | | Energies | | | | 4 HTMD | | | | 8 HTMD | | | |
|---|---|---|---|---|---|---|---|---|---|---|---|---|---|---|---|---|
| | RMSE | R | p-val | SD | RMSE | R | p-val | SD | RMSE | R | p-val | SD | RMSE | R | p-val | SD |
| Astex | 1.88 | 0.43 | 6.00E-02 | 1.41 | 1.77 | 0.39 | 9.00E-02 | 1.84 | 1.52 | 0.33 | 2.00E-01 | 1.03 | 1.74 | 0.08 | 7.00E-01 | 1.27 |
| CSAR | | | | | | | | | | | | | | | | |
| HIQ 1 | 2.01 | 0.71 | 2.00E-09 | 0.98 | 2.01 | 0.73 | 2.00E-10 | 0.81 | 2.01 | 0.74 | 1.00E-10 | 0.95 | 1.83 | 0.76 | 2.00E-11 | 0.91 |
| HIQ 2 | 1.59 | 0.75 | 7.00E-10 | 0.95 | 1.74 | 0.71 | 1.00E-08 | 0.9 | 1.62 | 0.72 | 4.00E-09 | 1.21 | 1.43 | 0.77 | 9.00E-11 | 1.07 |
| v.2014 | | | | | | | | | | | | | | | | |
| NMR | 1.61 | 0.4 | 2.00E-02 | 0.7 | 1.67 | 0.44 | 9.00E-03 | 1.11 | 1.42 | 0.62 | 1.00E-04 | 0.87 | 1.57 | 0.53 | 1.00E-03 | 1.48 |
| <=1 Å | 1.54 | 0.44 | 7.00E-05 | 0.88 | 1.62 | 0.39 | 6.00E-04 | 1.1 | 1.42 | 0.49 | 9.00E-06 | 0.79 | 1.61 | 0.41 | 3.00E-04 | 1.33 |
| <=1.5 Å | 1.7 | 0.26 | 3.00E-03 | 1.01 | 1.71 | 0.28 | 2.00E-03 | 1.13 | 1.51 | 0.34 | 1.00E-04 | 0.82 | 1.65 | 0.28 | 1.00E-03 | 1.25 |
| <=2 Å | 0.94 | 0.48 | 1.00E-12 | 0.5 | 1.33 | -0.05 | 5.00E-01 | 0.67 | 1.03 | 0.33 | 2.00E-06 | 0.52 | 0.95 | 0.4 | 5.00E-09 | 0.45 |
| FEP | 1.88 | 0.43 | 6.00E-02 | 1.41 | 1.77 | 0.39 | 9.00E-02 | 1.84 | 1.52 | 0.33 | 2.00E-01 | 1.03 | 1.74 | 0.08 | 7.00E-01 | 1.27 |

RMSE: Root Mean Square Error
R: Spearman's correlation coefficient
p-value: Significance of measured correlation
SD: Standard Deviation of the Linear Regression

**Table S5. T-test comparing the performance of RosENet to models trained during the feature ablation and network architecture analysis.** A pairwise T-test was used for computing t-values and p-values.

| | SqueezeNet | | | | ResNet | | |
|---|---|---|---|---|---|---|---|
| | 8 HTMD | 4 HTMD | Energies + 4 HTMD | Energies | 8 HTMD | 4 HTMD | Energies |
| t-value | -2.142 | -3.291 | -3.272 | -7.360 | -4.197 | -1.958 | -4.691 |
| p-value | 0.03 | 1e-03 | 1.1e-03 | 8.8e-13 | 3.2e-05 | 0.05 | 3.6e-06 |
